# Alveolar cell fate selection and lifelong maintenance of AT2 cells by FGF signaling

**DOI:** 10.1101/2022.01.17.476560

**Authors:** Douglas G. Brownfield, Alex Diaz de Arce, Elisa Ghelfi, Astrid Gillich, Tushar J. Desai, Mark A. Krasnow

**Affiliations:** Department of Biochemistry and Howard Hughes Medical Institute, Stanford University School of Medicine, Stanford, CA 94305-5307, USA; Molecular and Integrative Physiological Sciences Program, Harvard T.H. Chan School of Public Health, Boston, MA, USA; Department of Internal Medicine and Stem Cell Institute, Stanford University School of Medicine, Stanford, CA 94305, USA

**Keywords:** Lung development, alveolus formation, FGF signaling, cell fate selection, cell fate maintenance

## Abstract

The lung’s gas exchange surface comprises thin alveolar type 1 (AT1) cells and cuboidal surfactant-secreting AT2 cells that are corrupted in some of the most common and deadly diseases including adenocarcinoma, emphysema, and SARS/Covid-19. These cells arise from an embryonic progenitor whose development into an AT1 or AT2 cell is thought to be dictated by differential mechanical forces. Here we show the critical determinant is FGF signaling. FGF Receptor 2 (Fgfr2) is expressed in mouse progenitors then restricts to nascent AT2 cells and remains on throughout life. Its ligands are expressed in surrounding mesenchyme and can, in the absence of differential mechanical cues, induce purified, uncommitted E16.5 progenitors to form alveolus-like structures with intermingled AT2 and AT1 cells. FGF signaling directly and cell autonomously specifies AT2 fate; progenitors lacking *Fgfr2* in vitro and in vivo exclusively acquire AT1 fate. *Fgfr2* loss in AT2 cells perinatally results in reprogramming to AT1 fate, whereas loss or inhibition later in life immediately triggers AT2 apoptosis followed by a compensatory regenerative response. We propose Fgfr2 signaling directly selects AT2 fate during development, induces a cell non-autonomous secondary signal for AT1 fate, and stays on throughout life to continuously maintain healthy AT2 cells.

**One Sentence Summary:** FGF signaling induces and distinguishes the two cell types of the lung’s gas exchange surface, and the pathway remains on throughout life to maintain one that can be transformed into lung cancer or targeted in the deadly form of SARS/Covid-19.

## Introduction

Gas exchange occurs in alveoli, tiny terminal air sacs of the lung lined by two intermingled epithelial cell types: exquisitely thin alveolar type 1 (AT1) cells that provide the gas-exchange surface, and cuboidal AT2 cells that secrete surfactant to prevent alveolar collapse and acute respiratory failure^1^. Alveoli are also the site of some of the most significant but poorly understood and difficult to treat human diseases, including bronchopulmonary dysplasia in infants and COPD/emphysema^2^, pulmonary fibrosis^3^, lung adenocarcinoma^4, 5^, and viral infections including Severe Acute Respiratory Syndrome (SARS)^6, 7^ and the current pandemic of COVID-19^8^ in adults. Understanding how these key alveolar cell types arise during development and are maintained throughout life is critical for understanding and treating these diseases, and for guiding developmental, regenerative and tissue engineering approaches to create healthy alveoli and the surfactant that lines them.

Despite the extreme differences in structure and function of AT1 and AT2 cells, marker expression, lineage tracing, and clonal analysis in mice indicate that each arises directly from a common progenitor during embryonic development^5, 9^. This model is supported by single cell RNA sequencing studies of developmental intermediates that reconstructed the full, bifurcating gene expression program from bipotent progenitors at embryonic day 16.5 (e16.5) to either AT1 or AT2 cells over the next several days of fetal development^10–12^, although a recent genetic labeling study claims an earlier fate commitment^13^. Multiple developmental signaling pathways^14–18^ and transcription factors^19^ can influence alveolar structure, maturation or the balance of the two cell types^20–22^, but the key driver of alveolar fate selection and differentiation is thought to be mechanical forces^23^. The build up and increased movement of luminal fluid in late gestation is proposed to stretch and flatten progenitors into AT1 cells^18, 24, 25^, as stretch does to cultured AT2 cells to cause AT1 transdifferentiation in vitro^26^. Live imaging indicates that alveolar progenitors protected from the luminal mechanical forces during budding default to the AT2 cell program^18^.

Here we demonstrate that alveolar cell fate specification is in fact dictated by a classic growth factor signal, the same FGF signaling pathway that controls the earlier steps of airway budding and branching^22, 27–30^ and also initiates subsequent budding^18^. After branching, the receptor Fgfr2 remains on in alveolar progenitors and dynamically restricts to the AT2 lineage shortly before cellular differentiation, while its ligands Fgf7 and Fgf10 continue to be expressed by surrounding stromal cells. We show by the effects of the ligands on isolated e16.5 progenitors in cultures lacking budding and differential mechanical cues, and by genetic mosaic and pharmacological studies in culture and in vivo, that Fgfr2 signaling directly and cell autonomously specifies AT2 fate while inducing a secondary, non-autonomous signal that promotes AT1 fate of neighboring progenitors. This developmental FGF pathway remains on throughout life where it serves continuously and ubiquitously to maintain healthy AT2 cells^31, 32^, which if deprived of the FGF signal early in life reprogram to AT1 fate and later in life immediately undergo apoptosis.

## Results

### *Fgfr2* restricts to the AT2 lineage during alveolar differentiation

To identify signaling pathways that might control alveolar cell fate selection, we searched the single cell transcriptional program of mouse alveolar development^11^ for receptor genes enriched in either the AT1 or AT2 lineages. Of the five receptor genes showing at least two-fold-enrichment in the AT2 lineage (Fig. 1A; Fgfr2, Fzd8, Cd36, Cd74, and Ngfrap1), only Fgfr2 is known to be required for embryonic lung development^22, 30^. The other four either have no consequence (Fzd8), only adult lung phenotypes (Cd36, Cd74), or have not yet been examined (Ngfrap1). *Fgfr2* is expressed early and throughout the developing lung epithelium and its deletion abrogates airway branching^22, 29, 30^, so its later function has been more challenging to investigate^14, 31–33^. The single-cell RNA sequencing (scRNAseq) profiles demonstrated that the iiib isoform of *Fgfr2* is expressed in bipotent progenitors and maintained in the AT2 lineage but downregulated in the AT1 lineage (Fig. 1A, Supplementary Fig. 1C). Fgfr2 immunostaining during mouse alveolar development confirmed that Fgfr2 is expressed in bipotent progenitors (Fig. 1C), downregulated in nascent AT1 cells, and maintained in nascent AT2 cells as they bud into the surrounding mesenchyme and become histologically and molecularly distinct from AT1 cells (Fig. 1C; Supplementary Fig. 1A). Indeed, Fgfr2 is among the earliest known markers of AT2 fate selection.

**Figure 1.**
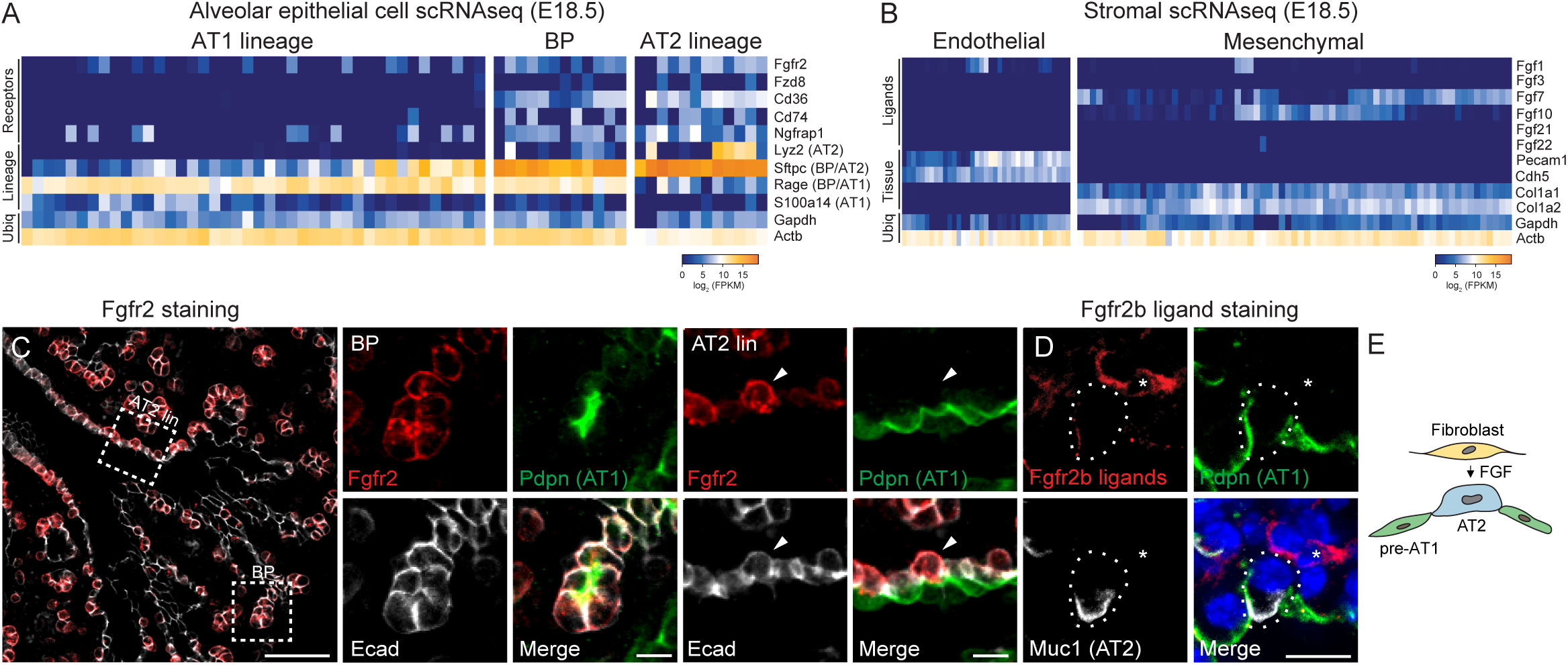
**Expression of Fgfr2 and its ligands during alveolar differentiation.** (A) Expression of *Fgfr2* and the four other receptor genes most selectively expressed in alveolar type 2 (AT2) cell lineage ("Receptors") from single-cell RNA sequencing (scRNAseq) analysis of distal (alveolar) epithelial cells from embryonic day 18.5 (e18.5) mouse lung. Cells (columns) are arranged in developmental pseudotime (BP, bipotent progenitors, center; AT1 lineage to left; AT2 lineage to right) determined as described^11^ by expression of alveolar lineage markers like the four shown ("Lineage") including two that restrict from BP to AT2 lineage (BP/AT2, *Sftpc*) or to AT1 lineage (BP/AT1, *Rage*). (Note due to the very high expression of *Sftpc* in bipotent progenitors, there is a temporal lag in extinction of the transcripts in newly-differentiating AT1 cells and thus some nascent AT1 cells co-express *Sftpc* and *Rage*.) Ubiq, ubiquitously-expressed control genes. Heat map, mRNA expression level. (B) Expression of Fgfr2b ligand genes in distal (alveolar) endothelium and mesenchyme from scRNAseq of e18.5 lung. (C) e17.5 lung immunostained for Fgfr2 (red), AT1 marker podoplanin (Pdpn, green), and epithelial marker E-cadherin (E-cad, white). Boxed regions, close-ups and split channels at right of bipotent progenitor (BP) and developing AT2 lineage cell (AT2 lin). Note Fgfr2 (red) remains on in developing AT2 cells (arrowhead) but downregulated in neighboring developing AT1 cells, and although AT1 marker (Pdpn) is still detected it will soon be downregulated as developing AT2 cell matures (see Supplementary Fig. 1A). Scale bars, 50µm (left panel), 10µm (close-ups). (D) e17.5 lung stained with the Fgfr2 (isoform iiib) ligand-binding domain fused to human IgG1 domain to show Fgfr2b ligands (red), and co-stained for Pdpn (green), and AT2 marker mucin1 (Muc1, white). Fgfr2b ligands are detected in nearby mesenchymal cells (asterisk) and diffusely around developing alveolar epithelial cells. Scale bar, 10µm. (E) Schematic showing inferred signaling from Fgf ligand-expressing mesenchymal cell (fibroblast) to nearby Fgfr2-expressing nascent AT2 cell during alveologenesis. pre-AT1, nascent AT1 cells that down-regulated Fgfr2.

To identify the relevant Fgfr2 ligands, we performed scRNAseq on the adjacent mesenchymal cells during alveolar differentiation. Two Fgfr2 (isoform iiib) ligands, *Fgf7* and *Fgf10,* were detected in mesenchymal cells (Fig. 1B), predominantly in the previously identified major subpopulation ("matrix fibroblast") marked by *Wnt2* expression (Supplementary Fig. 2A)^10^. Both *Fgf* genes are also expressed in lung mesenchyme earlier in development, when Fgf10 serves as the key ligand in Fgfr2-induced branching of the bronchial tree^22, 27^.

**Figure 2.**
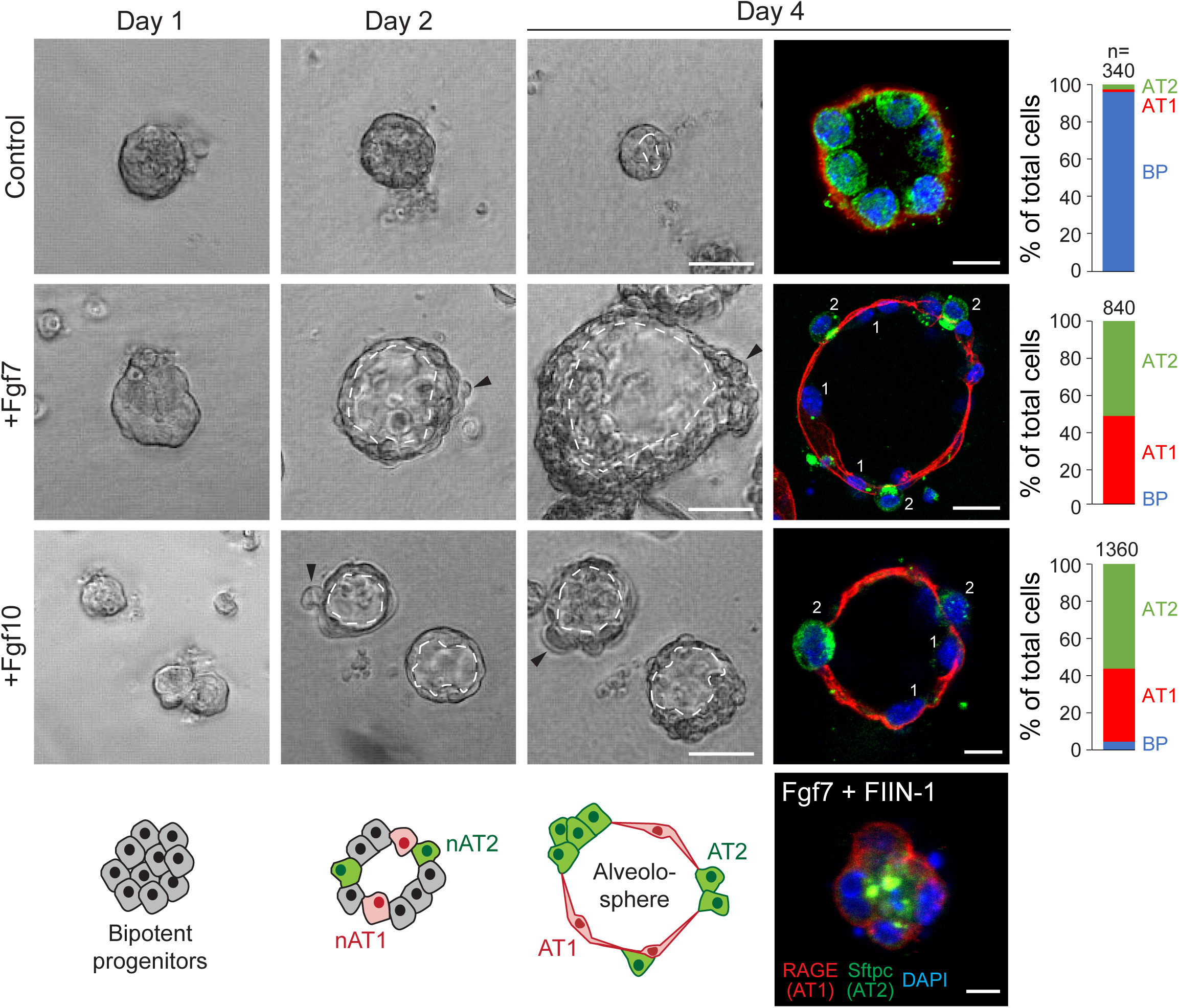
Effect of Fgfr2 ligands on purified alveolar epithelial progenitors in culture. Phase images (left panels) and immunostains (right) of epithelial progenitors purified from tips of e16.5 lungs and cultured in Matrigel for the period indicated with media alone (Control, top panels) or media supplemented every two days with Fgf7 (50 ng/ml, middle panels) or Fgf10 (100 ng/ml) and HSPG (100 ng/ml, lower panels). Note increase in luminal surface area (dashed lines) and basal extrusion (arrowheads) in cultures with Fgf ligands, and differentiation into intermingled cuboidal AT2 (Sftpcpos, green) and squamous AT1 cells (RAGE^pos^, red) at day 4 ("alveolospheres"). In control cultures lacking FGFs (top row) or treated both with Fgf7 and the Fgfr inhibitor FIIN-1 (bottom panel), cells remained bipotent progenitors throughout as shown by co-expression of both markers. nAT1, nascent AT1 cell; nAT2, nascent AT2 cell. Scale bars, 50µm (left panels and right +Fgf7 panel), 10µm (right top and +Fgf10 panels), and 10µm (Fgf7 + FIIN-1 panel). Co-treatment with both Fgf7 and Fgf10 (with HSPG) gave similar results as each treatment alone (data not shown). Quantification at right gives percent of each cell type (bipotent progenitor (BP), AT1, AT2) in culture at day 4 determined by immunostaining (n>340 cells scored for control, >840 cells for Fgf7 treated, and >1360 for Fgf10 treated in experimental triplicate). ***, p<0.0001 (Student’s t-test), BP abundance in control vs Fgf treatment conditions.

Visualization of Fgfr2 ligands with the Fgfr2 (isoform iiib) ligand-binding domain fused to human IgG1 domain showed ligand production in a subset of mesenchymal cells near budding AT2 cells (Fig. 1D, E; Supplementary Fig. 1B).

### Fgfr2 ligands induce alveolar differentiation and morphogenesis in culture

To explore the function of Fgfr2 signaling in alveolar development, we first examined the effect of Fgf7 and Fgf10 on purified epithelial progenitors isolated from the tips of e16.5 lungs, before AT1 and AT2 cells are detected. When the progenitors were cultured for up to 8 days in Matrigel in the absence of exogenous FGFs, they failed to develop into either AT1 or AT2 cells, remaining cuboidal and organizing into multicellular clusters around a lumen (Fig. 2; Supplementary Fig. 3C). By contrast, in the presence of Fgf7 (50 ng/ml), progenitor cells organized into large epithelial spheres containing differentiated cuboidal Sftpcpos Rage^neg^ AT2 cells intermingled with squamous Sftpc^neg^ Rage^pos^ Pdpn^pos^ AT1 cells, similar to the structures formed during alveologenesis in vivo beginning at ∼e17.5 (Fig. 2; Supplementary Fig. 3B).

Fgf10 induced formation of similar epithelial spheres composed of intermingled AT2 and AT1 cells when administered with heparan sulfate proteoglycans (HSPGs), a co-receptor that increases ligand accessibility^34, 35^ (Fig. 2). The effects of both Fgf7 and Fgf10 were abrogated in the presence of 10nM Fgfr inhibitor FIIN-1 (Fig. 2, Supplementary Fig. 3A). Live imaging of the cultures showed that Fgf7 stimulated continuous growth of the epithelial spheres (Supplementary Fig. 3F); however, no early cell budding like that proposed to protect progenitors from luminal mechanical forces during alveolar development in vivo^18^ was detected. Transient buds were occasionally observed but only after luminal expansion and AT1 flattening were apparent (Supplementary Fig. 3G). Likewise, the Arp2/3 inhibitor CK666 used to block protrusions and AT2 development in vivo^18^, reduced the transient buds but not Fgf7-induced AT1 and AT2 development in culture (Supplementary Fig. 3H). Thus, Fgfr2 signaling drives differentiation of alveolar progenitors and formation and growth of alveolus-like structures ("alveolospheres") with intermingled AT1 and AT2 cells, and it does so independent of extrinsic mechanical forces and without the early budding and associated structural features proposed to differentially protect progenitors from such forces. However, it is possible that extrinsic mechanical forces and budding *in vivo* might regulate some aspect of Fgf signaling or the developmental process that is dispensable in the culture system.

### Fgfr2 signaling directly selects AT2 cell fate

Addition of FGF ligands to the cultures triggered differentiation of progenitors into both AT1 and AT2 cells. To distinguish direct (cell autonomous) effects of FGF from secondary (cell non-autonomous) effects, we devised a strategy to enforce or block Fgfr2 signaling in random subsets of cultured progenitors targeted by a lentiviral vector. The vectors expressed either wild type Fgfr2 (Lenti-Fgfr2) or a dominant negative form lacking the cytoplasmic domain (Lenti-Fgfr2^DN^) along with a fluorescent reporter (GFP) to mark infected cells (Fig. 3A). We found that nearly every progenitor cell (95%) infected with virus expressing wild type Fgfr2 differentiated into an AT2 cell, and conversely infected cells expressing Fgfr2^DN^ almost always (88%) acquired AT1 fate (Fig. 3B,C); the rare cells that did not acquire the predominant fate remained undifferentiated (Fig. 3C, Supplementary Fig. 4A). Experiments with a control virus (AAV-nGFP) that expresses GFP but no Fgfr showed that randomly-infected progenitors have a roughly equal chance of acquiring AT1 or AT2 fate during culturing (Supplementary Fig. 4B).

**Figure 3.**
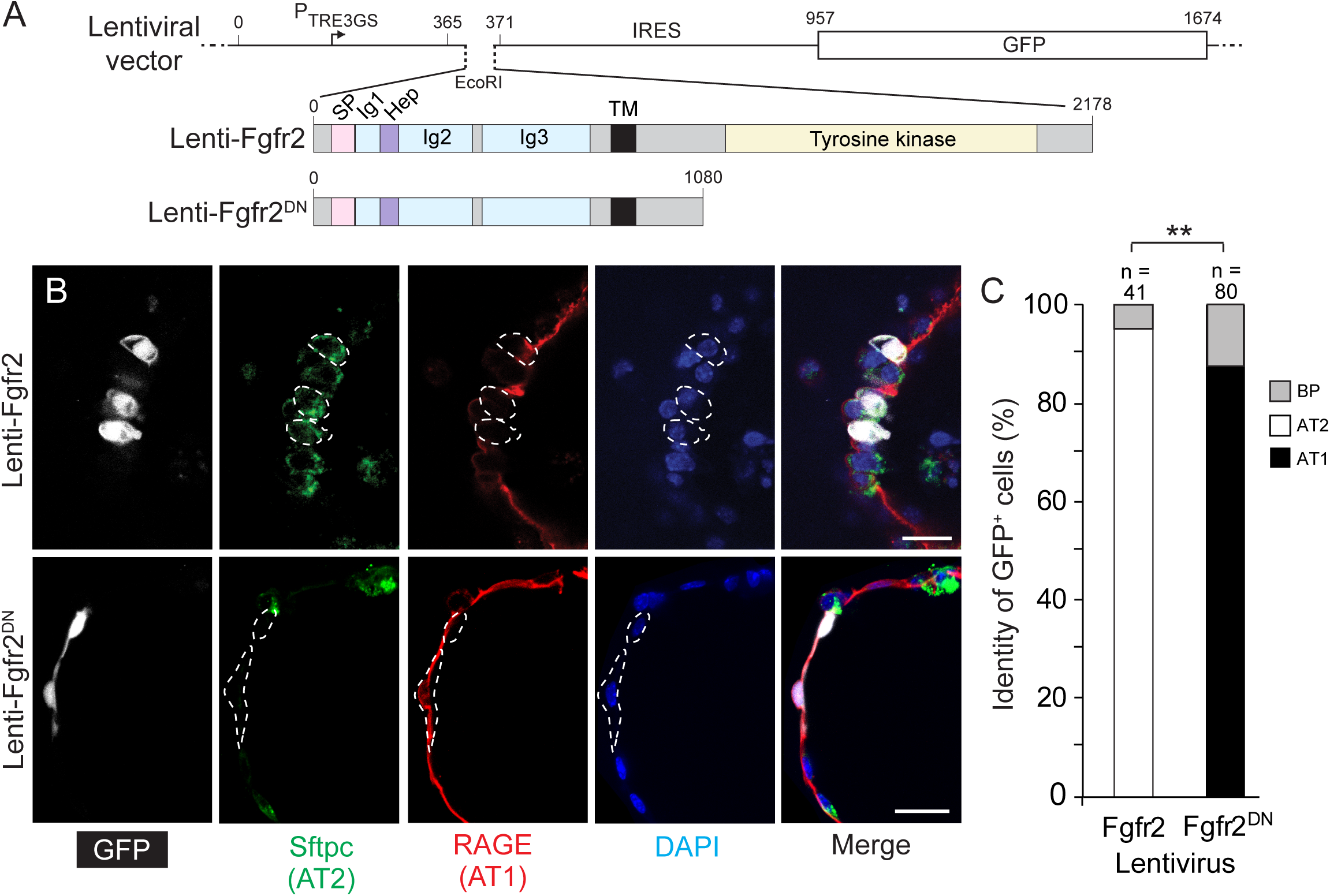
**Fgfr2 signaling controls AT2 fate selection cell autonomously in culture.** (A) Structures of lentiviral vectors co-expressing wild type Fgfr2 (Lenti-Fgfr2) or dominant negative Fgfr2 lacking the tyrosine kinase domain (Lenti-Fgfr2^DN^) and GFP. Numbers above vector indicate nucleotide position and above Fgfr2 structure indicate amino acid residue; colors highlight functional domains in Fgfr2. TRE, tetracycline responsive element; IRES, internal ribosome entry site; GFP, green fluorescent protein; SP, signal peptide; Ig, immunoglobulin domain; Hep, heparin sulfate binding domain; TM, transmembrane domain; DN, dominant negative. (B) e16.5 alveolar progenitors were mosaically infected (less than 1% of the cultured cells were infected) with the lentiviral vectors indicated at the time of cell plating in doxycycline-containing media (100 ng/ml) for 24 hours, then cultured with Fgf7 and doxycycline as in Fig. 2 for 4 days and immunostained for markers indicated. Note that GFP^pos^ cells in top panel with Fgfr2 signaling promoted cell autonomously by expression of wild type Fgfr2 became Sftpcpos RAGE^neg^ cuboidal AT2 cells, whereas GFP^pos^ cells in bottom panel with Fgfr2 signaling cell autonomously inhibited by expression of Fgfr2^DN^ became Sftpcneg RAGE^pos^ squamous AT1 cells. See Supplementary Figure 4A for images of rare infected cells that did not acquire the predominant fate. Scale bars, 20µm. (C) Quantification of B. Sftpcpos RAGE^pos^ cuboidal cells were scored as bipotent progenitors (BP). n= number of GFP^pos^ cells scored in 3 experiments. **, p<0.001 (chi-squared).

These results, together with the above result showing e16.5 distal progenitors cultured for over a week in the absence of FGF fail to differentiate into AT1 or AT2 cells (Fig. 2; Supplementary Fig. 3C), indicate that the fate of alveolar progenitors is not committed at e16.5, countering claims of an earlier fate commitment^13^ but consistent with prior lineage tracing^9^, clonal analysis^5^, and scRNAseq results^10–12^. We conclude that distal epithelial progenitors at e16.5 are indeed bipotent, having the capacity to differentiate into either AT1 or AT2 cells, and that AT2 fate is selected directly and cell autonomously by Fgfr2 signaling. The results also imply that the observed induction of AT1 fate by FGF addition to the cultures must be indirect (cell non-autonomous), presumably via a secondary signal produced by maturing AT2 cells and received by neighboring progenitors (see Discussion). AT1 differentiation cannot simply be the default fate because it too required addition of FGF ligands to the culture.

To determine if Fgfr2 signaling plays a similar role in alveolar cell fate selection in vivo, we used an *Nkx2.1-Cre* transgene with low Cre activity to mosaically delete *Fgfr2* in the developing lung epithelium, combined with a fluorescent Cre reporter allele (*Rosa26^mTmG^*) to mark the cells in which Cre was active (Fig. 4, Supplementary Fig. 4C). In control mice carrying a wild type *Fgfr2* allele in trans to the conditional *Fgfr2* allele (*Tg^Nkx2.1-Cre^;Fgfr2^fl/+^; Rosa26^mTmG/+^*), one third of distal cells with Cre reporter activity (36%, n> 970 GFP+ cells scored in 3 mice) acquired AT1 fate by the end of fetal life (PN0) and the rest acquired AT2 fate (64%) (Fig. 4A,B). By contrast, in animals with an *Fgfr2* deletion in trans to the conditional allele (*Tg^Nkx2.1Cre^;Fgfr2^fl/del^;Rosa26^mTmG/+^*), the targeted distal cells almost exclusively (91%, n≥460 GFP+ cells scored in 3 mice) acquired AT1 fate (Fig. 4A,B); the few cells that did not acquire AT1 fate had not yet lost Fgfr2 expression (Fig. 4C). We conclude that *Fgfr2* is the critical determinant of AT2 fate and is required cell autonomously for AT2 fate selection in vivo as well as in vitro, and that alveolar progenitors lacking Fgfr2 signaling acquire the alternative, AT1 fate.

**Figure 4.**
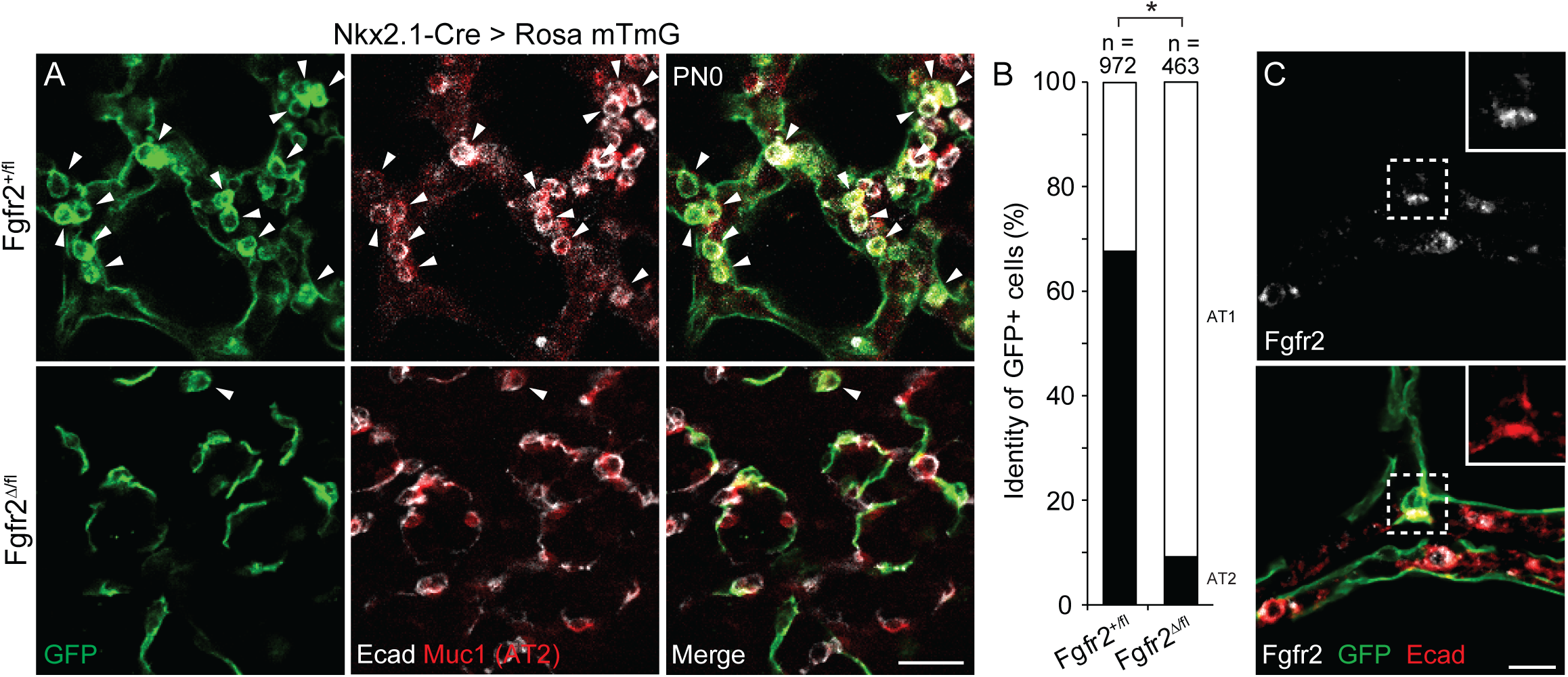
**Cell autonomous requirement of *Fgfr2* for AT2 fate selection in vivo.** (A) Alveolar region of lungs from *Nkx2.1-Cre; Rosa26-mTmG; Fgfr2^fl/+^* (upper panels) or *Nkx2.1-Cre; Rosa26-mTmG; Fgfr2^fl/delta^* (lower) mice at postnatal day 0 with Cre expressed in developing lung epithelial cells to delete conditional *Fgfr2* allele (*Fgfr2^fl^*) and activate a farnesylated GFP protein reporter, which targets all membranes including cytoplasmic vesicles (mTmG, green). Lungs were immunostained for GFP, E-cadherin (Ecad), and AT2 marker Muc1 as indicated. Note mosaic GFP expression showing Cre is active in some but not all alveolar epithelial cells, and that *Fgfr2* is required cell autonomously for AT2 cell development because GFP^pos^ control cells carrying wild type *Fgfr2* allele (*Fgfr2^fl/+^*, upper panels) became either cuboidal Muc1^pos^ AT2 cells (arrowheads) or squamous AT1 cells, whereas GFP^pos^ cells lacking *Fgfr2* (*Fgfr2^fl/delta^*, lower panels) almost exclusively became AT1 cells. Scale bar, 50µm. (B) Quantification of A. n= number of GFP^pos^ cells scored in 3 lungs. *, p< 0.005 (chi-squared). (C) Close up of a rare GFP^pos^ AT2 cell (boxed) from *Nkx2.1-Cre; Rosa26-mTmG; Fgfr2^fl/delta^* lung, as in bottom panels of A, stained for Fgfr2 and markers indicated. Note Fgfr2 has not yet been lost from GFP^pos^ cell, implying recent deletion of conditional allele *Fgfr2^fl^* and perdurance of Fgfr2 that promoted AT2 fate selection. Scale bar, 10µm.

### Fgfr2 signaling prevents AT2 cell reprogramming to AT1 fate during juvenile life

Curiously, this developmental signaling pathway remains on later in life, for months or even years after alveoli have acquired their canonical structure and function. *Fgfr2* continues to be selectively expressed in AT2 cells (Fig. 5A,B,D; Supplementary Fig. 5), *Fgf7* and *Fgf10* continue to be expressed in *Wnt2*-expressing alveolar fibroblasts and at lower levels in myofibroblasts (Fig. 5E-I, Supplementary Fig. 2B,C), and the Fgfr2 signaling pathway remains active in mature AT2 cells as detected by phosphorylated MAP kinase (Fig. 5C,D) and expression of Fgfr2-induced genes such as *Spry2* (Fig. 5A). To investigate the role of this selective and persistent Fgfr2 signaling in AT2 cells, we first used a *Cre* knock-in allele at the endogenous *Lysozyme2* (*Lyz2*) locus, which turns on in some AT2 cells early in postnatal life and is progressively activated later in other AT2 cells^5^ (see below), to conditionally delete Fgfr2 in AT2 cells at different ages. There was a dramatic effect of Fgfr2 deletion from AT2 cells during juvenile life, and an equally significant but entirely different effect in adults.

**Figure 5.**
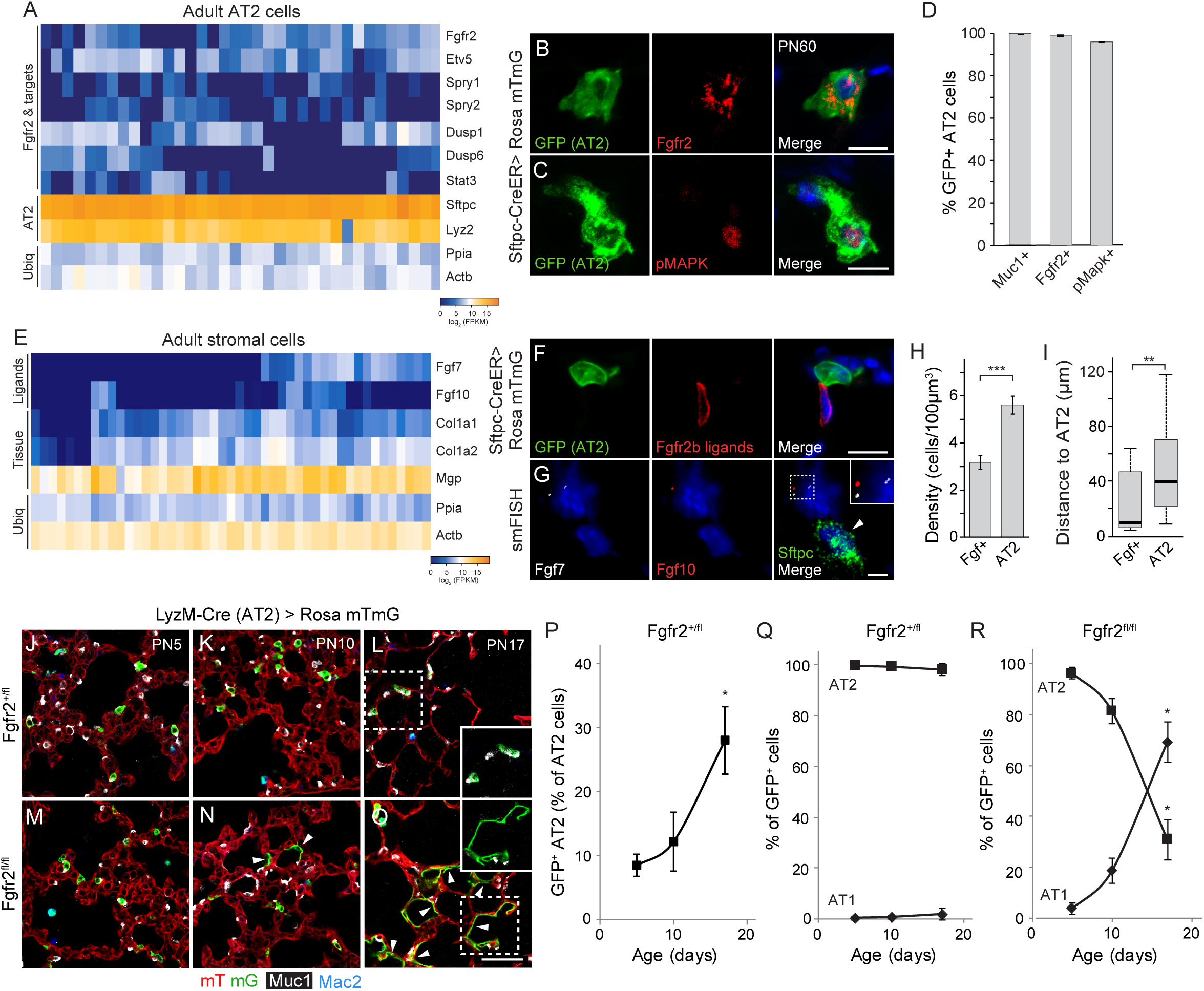
Fgfr2 pathway remains on after development and prevents AT2 reprogramming to AT1 fate in juvenile period. (A) Expression of *Fgfr2* and downstream transcription factor *Etv5* and target genes in adult AT2 cells by scRNAseq indicates pathway remains on throughout life in AT2 cells. AT2, AT cell markers; Ubiq, ubiquitously-expressed control genes. (B,C) Close-up of AT2 cells in adult (2 months) *Sftpc-CreER; Rosa26-mTmG* mouse induced with tamoxifen one week prior to sacrifice to label with mGFP and co-stained for Fgfr2 (B) or phosphorylated MAP kinase (pMAPK) to show pathway activity (C) and counterstained with nuclear stain DAPI (blue). Scale bars, 10µm. (D) Quantification of B, C. Nearly all labeled adult AT2 cells (GFP^+^) express an AT2 marker (Muc1) and Fgfr2, and show Fgfr2 signaling activity (pMapk^+^). n=300 GFP^pos^ cells scored in 3 animals. (E) Expression of *Fgfr2* ligands in adult AT2 stromal cells by scRNAseq. (F) Close-up of AT2 cell in adult (2 months) *Sftpc-CreER; Rosa26-mTmG* mouse induced as above (panels B,C) to label AT2 cells with mGFP and co-stained for Fgfr2 (isoform iiib) ligand-binding domain to show Fgfr2b ligands (red) and counterstained with DAPI. This provides an overview of the spatial relationships between FGF sending and receiving cells but does not identify the FGF-expressing cells, which is identified by scRNA-seq in Supplementary Fig. 2). Scale bar, 10µm. (G) Single molecule fluorescence in situ hybridization (smFISH) for *Fgf7*, *Fgf10*, and *Sftpc* in adult (PN60) mouse lung. Note cell neighboring AT2 cell that co-expresses *Fgf7* (white dots) and *Fgf10* (red dots) (enlarged in inset). Blue, DAPI (nuclear counterstain. Scale bar, 5µm. (H,I) Quantification of G showing abundance of *Fgf*-expressing and AT2 cells (H, *** p-value < 0.0001, Student’s t-test ) and distance between them (I, ** p-value <0.001, Mann Whitney test). n=182 AT2 and 103 Fgf^+^ cells scored in 4 animals. Because individual alveolar cells have complex structures we cannot accurately determine the number of AT2 cells directly contacted by an FGF-expressing cell; these cell density and distance measurements provide a sense of the availability of Fgf ligand and how many AT2 cells it supports, as well as how far each Fgf-expressing cell would need to extend (or the ligand diffuse) to support AT2 cells. (J-O) Alveolar region of *LyzM-Cre; Rosa26-mTmG; Fgfr2^fl/+^* (J-L) or *LyzM-Cre; Rosa26-mTmG; Fgfr2^fl/fl^* (M-O) mice at indicated postnatal day immunostained for Cre reporter mTmG, AT2 marker Muc1, and macrophage marker Mac2 to distinguish alveolar macrophages from AT2 cells. *LyzM-Cre* becomes active in AT2 cells in early postnatal life and activates GFP reporter and deletes conditional *Fgfr2* allele (*Fgfr2^fl^*). Note GFP^pos^ control AT2 cells carrying a wild type *Fgfr2* allele (*Fgfr2^fl/+^*, J-L) remain as AT2 cells (boxed inset in L), whereas GFP^pos^ cells lacking *Fgfr2* (*Fgfr2^fl/fl^*, M-O) convert to AT1 cells during this period (arrowheads and boxed inset in O). Scale bar, 50µm. (P-R) Quantification of J-O showing induction of Cre activity in AT2 cells in early postnatal life (P) and identities of GFP^pos^ cells in control (Q) and following *Fgfr2* loss (R). (P-R) n≥ 300 GFP^pos^ cells scored in 3 animals for each timepoint and condition (*p-value ≤ 0.01, Student’s t-test, time series end vs. start point).

Lyz2-Cre activity is first detectable in rare AT2 cells at PN5, as shown by the Cre reporter (Fig. 5J), and then gradually becomes active in additional AT2 cells (Fig. 5P, 6A (left panel). In juvenile control mice carrying an *Fgfr2* wild type allele (*Lyz2^Cre/+^;Fgfr2^fl/+^;Rosa26^mTmG/+^*), nearly all (>96%) AT2 cells expressing the Cre reporter retained their AT2 identity (Fig. 5J-L,Q), as previously observed for *Fgfr2^+/+^* AT2 cells^5^. By contrast, in juvenile mice homozygous for the *Fgfr2* conditional allele (*Lyz2^cre/+^Fgfr2^fl/fl^;Rosa26^mTmG/+^*), there was a rapid decline in the proportion of reporter-expressing cells with AT2 identity, from 96% at PN5 to 31% at PN17, and a concurrent increase in labeled cells that had acquired AT1 fate, from 3.8% at PN5 to 69% at PN17 (Fig. 5M-O,R).

**Figure 6.**
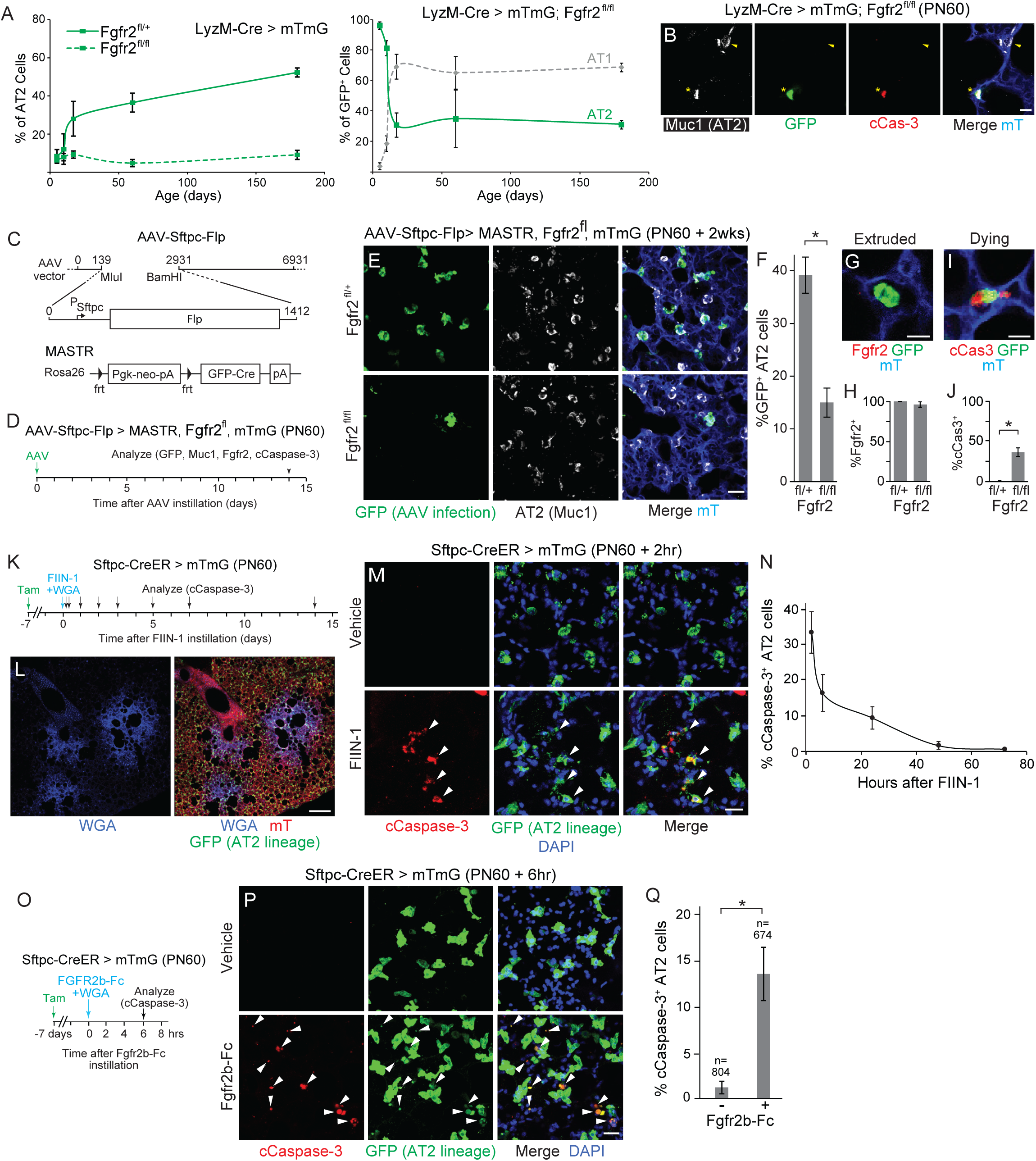
*Fgfr2* is required for survival of adult AT2 cells. (A) (Left panel) Time course showing the proportion of GFP^pos^ AT2 (Muc1^+^) cells in either control (*LyzM-Cre; Rosa26-mTmG Fgfr2^fl/+^*) or *Fgfr2* deleted (*LyzM-Cre; Rosa26-mTmG; Fgfr2^fl/fl^*) lungs at indicated postnatal times (control plot is extension of Fig. 5P). Note there is no further accumulation of GFP^pos^ AT2 cells in *Fgfr2* deleted lungs (dashed line) later in life (after day 20), despite continued LyzM-Cre activity and AT2 cell labeling apparent in the control lungs (solid line), suggesting *Fgfr2* deletion during adult life results in loss of the labeled cell. n=3955 AT2 cells scored in 3 animals per genotype. (Right panel) Time course showing identities of GFP^pos^ cells following *Fgfr2* loss in *LyzM-Cre; Rosa26-mTmG; Fgfr2^fl/fl^* mice at indicated postnatal times (extension of Fig. 5R plot). After the rapid increase in AT1 cells due to AT2 reprogramming in the juvenile period (through day 20), there was no further accumulation of GFP^pos^ AT1 or AT2 cells later in life despite continued LyzM-Cre activity and AT2 cell labeling, suggesting *Fgfr2* deletion during adult life results in loss of the labeled cell. n≥ 300 GFP^pos^ cells scored in 3 animals for each condition and timepoint. (B) Close-up of alveolar region in adult (2 months) *LyzM-Cre; Rosa26-mTmG; Fgfr2^fl/fl^* mouse stained for the indicated markers and cleaved Caspase-3 (cCaspase-3) to detect apoptotic cells. Note GFP^pos^ AT2 cell deleted for *Fgfr2* undergoing apoptosis (asterisk) but not the unlabeled control (*Fgfr2^+^)* AT2 cell nearby (arrowhead). Scale bar, 10µm. (C-F) Using MASTR system to test effect of conditional, complete deletion of *Fgfr2* in AT2 cells in adult mouse lung. (C) Structure of recombinant AAV-Sftpc-Flp virus (top) that expresses Flp recombinase from AT2-specific *Sftpc* promoter (top), and MASTR allele which, following Flp-mediated deletion of *Pgk-neo-pA* cassette using frt sites (triangles), constitutively expresses a GFP-Cre recombinase fusion protein. (D) Experimental scheme. Two weeks after AAV-Sftpc-Flp instillation into lungs of adult (PN60) *Fgfr2^fl/fl^; Rosa26^mTmG/MASTR^* mice to induce constitutive GFP-Cre expression and complete deletion of *Fgfr2^fl/fl^* in AT2 cells, lungs were harvested and stained for GFP, Muc1, Fgfr2, and cCaspase-3. (E) Close-up of alveolar region of lungs from heterozygous *Fgfr2^fl/+^* control (top panels) or *Fgfr2*^fl/fl^ (bottom) mice treated as in D. Note reduction of GFP-Cre expressing (GFP^pos^, green) AT2 cells (Muc1^pos^) in bottom panel, indicating complete deletion of *Fgfr2* in AT2 cells leads to AT2 cell loss. Scale bar, 25µm. (F) Quantification of AT2 cell loss in E from MASTR-mediated AT2 cell deletion of *Fgfr2* (*Fgfr2^fl/fl^*). Note 64% fewer GFP^pos^ AT2 (Muc1^pos^) cells in the experimental (*Fgfr2^fl/fl^*) vs control (*Fgfr2^fl/+^*) condition. n>890 Muc1^pos^ cells scored for each condition in 3 experiments. (G, H) Quantification (H) of Fgfr2 immunostaining (n≥100 GFP^pos^ cells scored for each condition in three experiments) shows that nearly all (96%) of the GFP^pos^ AT2 cells that remain after MASTR-mediated deletion of *Fgfr2^fl/fl^* (E, F) have not fully lost Fgfr2. Micrograph (G) shows close-up of one of the rare GFP^pos^ Fgfr2^neg^ AT2 cells detected, which like this one had each been extruded into alveolar space and showed rounded appearance. Scale bar, 10µm. (I, J) Quantification (J) of cCaspase-3 immunostaining (n≥470 GFP^pos^ cells scored for each condition) shows that 36% of GFP^pos^ AT2 cells that remain after MASTR-mediated deletion of *Fgfr2^fl/fl^* are cCaspase-3^pos^, indicating they have initiated apoptosis. Micrograph (I) shows close-up of a cCaspase-3^pos^ GFP^pos^ AT2 cell that has initiated apoptosis. Scale bar, 10µm. (K-N) Effect of acute inhibition of Fgfr2 signaling in AT2 cells by Fgfr inhibitor FIIN-1. (K) Experimental scheme for co-instillation of FIIN-1 and fluorescent marker WGA-405 into lungs of adult (PN60) *S*ftpcCreER/^+^; *Rosa26^mTmG/+^* mice treated with tamoxifen (Tam) to label mature AT2 cells and analyzed over two weeks. Scale bar, 200µm. (L) Lung regions exposed to instilled FIIN-1 shown by WGA-405 labeling (blue, left panel) and shown merged (right panel) with mTmG fluorescence with AT2 lineage cells labeled with GFP and all other cells labeled with tdTomato (mT). Scale bar, 250µm. (M) Close-up of WGA-405-labeled alveolar regions exposed for 2 hours to vehicle alone (top panels) or FIIN-1 (bottom). Note AT2 cells (GFP^pos^, green) exposed to FIIN-1 undergoing apoptosis (cCaspase-3^pos^, arrowheads) but not control AT2 exposed to vehicle alone. Scale bar, 20µm. (N) Quantification of E showing time course of AT2 apoptosis following FIIN-1 instillation. In vehicle control, <2.5% of AT2 cells were cCaspase-3^pos^ (n≥650 cells scored for each condition in 3 experiments). No conversion of labeled AT2 cells to AT1 identity was detected during this period (n≥390 cells scored for each condition in 3 experiments) (see also Suppl. Fig. 7). (O-Q) Effect of acute inhibition of Fgfr2 signaling in AT2 cells by instillation of a recombinant mouse Fgfr2iiib-Fc protein. (O) Experimental scheme. Seven days after tamoxifen (Tam) treatment of adult (PN60) SftpcCreER/+*; Rosa26^mTmG/+^* mice to GFP label AT2 cells, Fgfr2iiib-Fc protein (or vehicle as control) and fluorescent marker WGA-405 were co-instilled into the lungs to block its ligands in marked areas, and 6 hours later lungs were stained for GFP and apoptosis marker cCaspase-3 and counterstained with DAPI. (P) Close-ups of WGA-labeled alveolar regions from vehicle control (top panels) and Fgfr2iiib-Fc (bottom) instilled lungs. Scale bar, 25µm. (Q) Quantification of L. n, number of cells scored in 6 experiments. *, p-value <0.01, (Student’s t-test) (panels F, J, Q)

Deletion of *Fgfr2* with an *Sftpc-CreER* at PN0 can also promote AT1 fate^36^. A similar result obtained when Fgfr2 signaling was inhibited by addition of FIIN-1 to newly formed AT1 and AT2 cells generated from bipotent progenitors cells in culture, resulting in an increase in AT1 and reduction in AT2 cells (Supplementary Fig. 3D,E). We conclude that Fgfr2 signaling is necessary to maintain AT2 fate in newly formed AT2 cells during juvenile life, and that loss of Fgfr2 signaling during this period results in rapid and robust reprogramming to AT1 fate.

### *Fgfr2* signaling continuously prevents AT2 apoptosis during adult life

After the juvenile period (PN17) in the above experiment, there was no further increase in AT2-lineage-labeled cells with AT1 fate (Fig. 6A, right panel; Supplementary Fig. 6), indicating that AT2 cells that lose *Fgfr2* in adult life do not reprogram to AT1 fate. There was also no further accumulation of lineage-labeled AT2 cells during this period (Fig. 6A, right panel), suggesting that *Fgfr2* deletion in adult AT2 cells results in their loss. Indeed, we occasionally detected expression of cleaved Caspase-3 in such cells, indicating they had initiated apoptosis (Fig. 6B). Recently, others have noted a variable reduction in AT2 cell markers or abundance and impaired recovery from severe lung injury when one or more *Fgfr* genes were conditionally deleted^31–33^, attributing the deficits to a reduction in stem cells^31^ or their proliferation^32, 33^. However, another recent study reported *Fgfr2* in AT2 cells is entirely dispensable during homeostasis^36^. To more precisely determine the cellular and molecular consequences of acute abrogation of FGF signaling in adult AT2 cells, we first tried a tamoxifen-inducible Cre recombinase *(S*ftpcCreER/+; *Fgfr2^del/fl^; Rosa26^mTmG/+^*) but found that removal of *Fgfr2* occurred but was inefficient under these conditions (Fig. 4C) (see below and Methods). We therefore developed three additional approaches, which revealed that Fgfr2 signaling is ubiquitously and continuously required for AT2 cell maintenance throughout adult life, and even brief deprivation of signaling immediately initiates apoptosis.

Two important limitations of the conditional *Fgfr2* deletion experiment noted above were that the gene was inefficiently deleted, and even when the gene was deleted the protein persisted (perdured), limitations that could have contributed to the modest or entire lack of effect observed in other studies^32, 36^. To overcome the former limitation, we designed a combinatorial genetic approach using the MASTR transgene^37^ (Fig. 6C), which following Flp-mediated recombination of the transgene provides constitutive and high level Cre expression to ensure recombination of all Cre-dependent alleles in the cell, such as *Fgfr2^fl^* and *Rosa26^mTmG^*. We used the MASTR allele in conjunction with an AAV virus we engineered to express Flp recombinase specifically in AT2-cells using an *Sftpc* promoter element (Fig. 6C,D); when the virus was instilled intratracheally into adult control mice (*Fgfr2^+/fl^; Rosa26^mTmG/MASTR^*), specific expression of Cre-dependent reporter genes was observed in AT2 cells (>98% of GFP^+^ cells, n = 3 lungs, >168 GFP^+^ cells counted per lung (data not shown) at 2 weeks (Fig. 6E), similar to the AT2 specificity observed in prior studies with this approach^38–40^. When the virus was instead instilled into *Fgfr2^fl/fl^; Rosa26^mTmG/MASTR^* mice, 60% fewer GFP^+^ AT2 cells were observed (Fig. 6E,F); of the GFP^+^ AT2 cells that remained, nearly all (96%) were found to have retained Fgfr2 protein due to perdurance (Fig. 6H), and the rare GFP^+^ Fgfr2^-^ AT2 cells detected (4 of 96 scored GFP^+^ AT2 cells from 3 mice) had been extruded into the alveolar space and had a rounded morphology (Fig. 6G). This result indicates that adult AT2 cells broadly and uniformly undergo apoptosis following removal of Fgfr2. Apoptosis may even initiate with only partial reduction in Fgfr2 levels because one-third (36%) of the remaining GFP^+^ AT2 cells in this experiment, nearly all of which expressed detectable levels of Fgfr2 (Fig. 6H), were positive for cleaved-Caspase-3 (Fig. 6I,J).

To overcome the problem of Fgfr2 perdurance in AT2 cells and corroborate these results, we employed two other approaches. In a classical pharmacologic approach (Fig. 6K), we instilled Fgfr inhibitor FIIN-1 into the lungs of adult (PN60) SftpcCreER/+*; Rosa26^mTmG/+^* mice treated with tamoxifen two weeks earlier to label mature AT2 cells. A fluorescent marker (WGA-405) was included in the instillation to identify regions of FIIN-1 exposure (Fig. 6L). As early as two hours after the instillation, inhibition of Fgfr signaling triggered widespread apoptosis of AT2 cells in the FIIN-1 exposed (WGA-405 labeled) regions (Fig. 6M,N), with no conversion to AT1 identity (Supplementary Fig. 7). To ensure this effect was attributable to Fgfr2 inhibition, we also tested a soluble recombinant Fgfr2b-Fc protein that binds its cognate ligands (see Fig. 1), preventing their engagement and activation of the receptor (Fig. 6O). With this inhibitor too, there was substantial induction of AT2 cell apoptosis in the exposed regions shortly (6 hours) after instillation (Fig. 6P,Q). Thus, acute inhibition of Fgfr2 or its ligands in the adult lung rapidly and robustly induces AT2 cell apoptosis.

Akt (protein kinase B), a serine-threonine kinase, functions downstream of Fgfr signaling to mediate cell survival in other contexts^41–43^. To determine if Akt signaling functions downstream of Fgfr2 signaling to mediate AT2 cell survival in adult mouse lung, we activated Akt signaling by intraperitoneal injection of the small molecule Akt activator SC79 30 min prior to Fgfr2b inhibition by Fgfr2b-Fc installation. Activation of Akt almost completely abrogated AT2 apoptosis, as assayed by cleaved Caspase 3 (Supplementary Fig. 8). This suggests that Fgfr2 signaling promotes survival of adult AT2 cells through activation of Akt.

### Loss of AT2 cells induces a robust regenerative response

Despite the profound effects of Fgfr2 loss or inhibition on AT2 cells in juvenile and adult life, curiously alveolar structure and the total number and proportion of AT2 cells were not grossly altered (Fig. 7A, B), obscuring its ubiquitous and constant requirement for AT2 cell maintenance throughout life. This is because apoptosis (or reprogramming) of targeted AT2 cells is almost completely compensated by proliferation of untargeted cells. This was evident in the Lyz2-Cre mice (*Lyz2^cre/+^Fgfr2^fl/fl^;Rosa26^mTmG/+^*), where the population of untargeted AT2 cells (lacking the Cre-dependent lineage label) in PN60 mice was increased by 52% relative to that in the Fgfr2+ control *(Lyz2^Cre/+^;Fgfr2^fl/+^;Rosa26^mTmG/+^*), maintaining the normal density of AT2 cells (∼5 per 100 um^3^) and alveolar morphology despite the substantial loss of lineage-labeled AT2 cells (reduced from ∼36% of AT2 cells in control *Lyz2^Cre/+^;Fgfr2^fl/+^;Rosa26^mTmG/+^* mice to 2.8% in *Lyz2^cre/+^Fgfr2^fl/fl^;Rosa26^mTmG/+^* mice) (Fig. 7C). The proliferative response of remaining AT2 cells following the loss of Fgfr2 activity in neighboring cells was apparent when loss was initiated by pharmacologic inhibition (Fig. 7E-G) or even following the inefficient deletion of *Fgfr2* in *S*ftpcCreER/*^+^*; *Fgfr2^del/fl^* mice (Fig. 7D). The restorative response begins quickly because compensatory AT2 cell proliferation was detected by Ki67 immunostaining and EdU incorporation within hours of instillation of the Fgfr inhibitor FIIN-1 and initiation of AT2 apoptosis (Fig. 7F,G), and it continued for a week until alveolar structure was restored (Fig. 7G,H; Supplementary Fig. 9). The proliferating AT2 cells carrying out the restorative response (with Fgfr2 signaling intact because they were not from a FIIN-1-exposed area or because FIIN-1 had since cleared) are executing the canonical stem cell function of AT2 cells^5, 44, 45^, replacing lost AT2 cells and any AT1 cells lost along with them in the FIIN-1-exposed areas (Fig. 7H).

**Figure 7.**
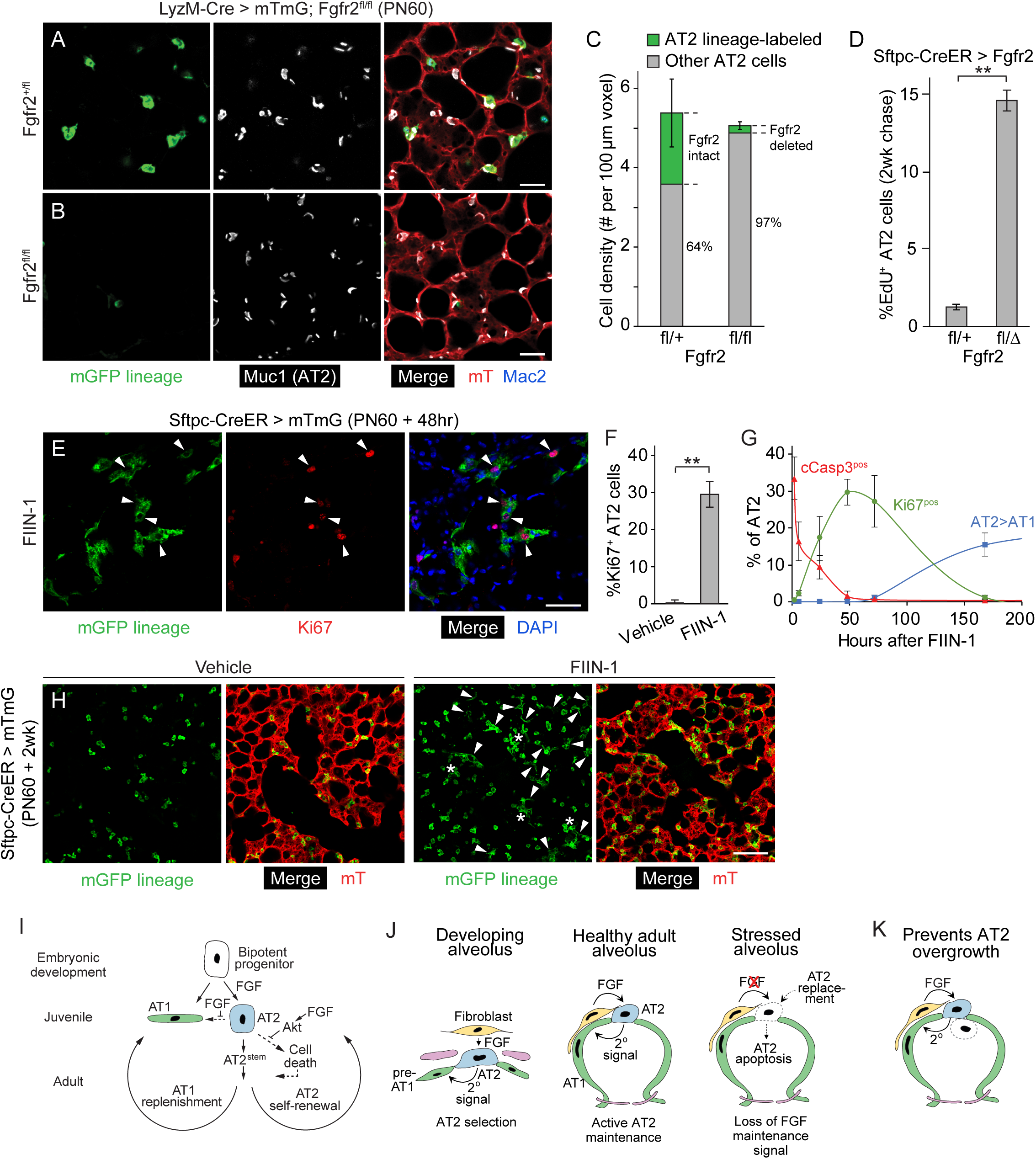
Robust regeneration following focal AT2 loss. (A,B) Alveolar regions of adult (PN60) control Fgfr2^+^ *LyzM-Cre; Rosa26-mTmG; Fgfr2^fl/+^* (A) and AT2 cell conditional deletion *LyzM-Cre; Rosa26-mTmG; Fgfr2^fl/fl^* (B) mice immunostained for markers indicated. Note loss of GFP lineage-labeled Muc1^pos^ AT2 cells in B but preservation of alveolar architecture. Mac2, macrophage marker to distinguish alveolar macrophages from labeled AT2 cells. Scale bars, 50µm. (C) Quantification of A and B. Note density of AT2 cells is maintained in AT2 cell conditional Fgfr2 deletion *(Fgfr2^fl/fl^*) condition by compensatory expansion of unlabeled AT2 cells (with unrecombined Fgfr^fl^ alleles) replacing lost lineage-labeled AT2 cells. n>300 cells scored for each condition in 6 animals; p=0.65 (not significant, Student’s t-test) for difference in cell density. (D) Quantification of AT2 cell proliferation in control adult (PN60) *Fgfr2^+^ Sftpc-CreER; Rosa26-mTmG; Fgfr2^fl/+^* and AT2 cell conditional *Fgfr2* deletion *Sftpc-CreER; Rosa26-mTmG; Fgfr2^fl/delta^* lungs by EdU labeling for 2 weeks following tamoxifen induction. Note conditional loss of *Fgfr2* induces compensatory increase in AT2 proliferation. n>400 AT2 cells scored for each condition in 6 lungs. **, p<0.001 (Student’s t-test). (E) Alveolar region of adult (PN60) *Sftpc-CreER; Rosa26-mTmG* induced with tamoxifen (PN60) to GFP label AT2 cells and instilled with Fgfr inhibitor FIIN-1 and stained 48 hours later for proliferation marker Ki67 and markers indicated. Note proliferating (Ki67^pos^) AT2 (GFP^pos^) cells (arrowheads). Scale bar, 50µm. (F) Quantification of E. n>800 AT2 cells scored for each condition in 6 animals. **, p<0.005 (Student’s t-test). (G) Time course of AT2 apoptosis (cCasp3^pos^) followed by compensatory AT2 proliferation (Ki67^pos^) and conversion to AT1 fate (Pdpn^pos^) after instillation of FIIN-1 in adult (PN60) SftpcCreER/+; Rosa*26^mTmG/+^* lungs as in Fig 6C-F and subsequent staining for AT2 lineage label mGFP, cCasp3, Ki67, and Pdpn. AT2 cells were scored only in instilled (WGA^pos^) alveolar regions; uninstilled regions and control vehicle-instilled lungs showed no changes. n>300 AT2 cells scored for each stain at each time point in 21 animals. (H) Vehicle (left) and FIIN-1-treated (right) alveolar regions as in E-G two weeks after instillation. Note clusters of lineage-labeled AT2 cells (asterisks) and conversion to AT1 fate (arrowheads) in FIIN-1-treated lung with overall restoration of alveolar structure. Scale bar, 100µm. (I) Sequential roles of Fgf/Fgfr2 pathway in alveolar development and maintenance: AT2 fate selection and 2° induction (see below) of AT1 fate (embryo), AT2 fate consolidation and prevention of reprogramming to AT1 fate (juvenile), and AT2 survival (adult), the latter mediated by Akt. AT2 cell death triggers a robust regenerative response presumably mediated by alveolar stem cells (AT2^stem^). (J) Source and dynamics of FGF signaling in developing, healthy, and stressed alveoli. After FGF selection of AT2 fate in development (left panel), a 2° signal induces other progenitors to AT1 fate. FGF signaling remains activate in adult to maintain AT2 cells (middle panel). Loss of FGF (red x) in stressed alveolus (right panel) results in AT2 cell death and rapid replacement. (K) FGF signaling in the adult alveolus might prevent AT2 overgrowth and tumor initiation by depriving daughter cells (dashed outline) that move away from the FGF source (e.g., into the lumen) of a critical survival cue.

## Discussion

The results identify FGF signaling, not differential mechanical forces, as the critical factor that induces and controls alveolar epithelial fate selection. *Fgfr2* is expressed in bipotent alveolar progenitors but rapidly restricts to nascent AT2 cells and remains on exclusively in the AT2 lineage, while its ligands Fgf7 and Fgf10 are expressed in surrounding mesenchyme. Addition of either ligand to cultured progenitors induced formation of alveolus-like structures with intermingled AT2 and AT1 cells, in the absence of any extrinsic forces or cell budding that might protect some progenitors from such forces. In mosaic cultures, randomly selected progenitors with constitutive *Fgfr2* expression exclusively acquired AT2 fate whereas those with the Fgfr2 pathway inhibited exclusively acquired AT1 fate, and a similar result obtained for *Fgfr2^-^* progenitors in mosaic alveoli in vivo. The results demonstrate that Fgfr2 signaling serves as a developmental switch in alveolar development, inducing bipotent progenitors to AT2 fate (Fig. 7I); progenitors with no or low Fgfr2 signaling acquire AT1 fate through a secondary, cell non-autonomous signal provided by nascent AT2 cells. Recently, we identified a secondary signal (Notch pathway) required for AT1 fate selection (Gillich et al, submitted); we suspect this secondary signal could be enhanced by (or synergize with) some subsequent local mechanical change, which would integrate our findings with the longstanding models implicating mechanical forces in alveolar fate acquisition^18, 23–25, 46^.

The FGF developmental pathway remains on in AT2 cells throughout life. Loss of Fgfr2 signaling during the juvenile period results in AT2 reprogramming to AT1 fate, perhaps by direct reprogramming without proliferation^47^ or by transient reversion to bipotent progenitor identity (Supplementary Fig. 3D,E). Loss or reduction of Fgfr2 signaling during adult life, even briefly at any time or position in the lung, triggers rapid AT2 apoptosis, followed by a robust regenerative response (Fig. 7I). The anti-apoptotic effect of Fgfr2 signaling is transduced through Akt. Thus, the Fgfr2 pathway directly selects AT2 cells during development and then ubiquitously and continuously maintains them throughout life.

The Fgfr2 pathway also plays a critical role in the first six days of lung development, inducing and patterning airway branching^22, 27, 29, 30, 48^. The signaling pathway is thus reused throughout the life of the animal but with changing roles and cellular consequences at each stage. As the lung forms in the embryo (e10-e16), Fgfr2 signaling provides mitogenic and motogenic functions to pattern airway epithelium budding and branching. During late fetal life (e16-e19) its budding function may continue as it selects alveolar cell fates by directly inducing AT2 differentiation and triggering a secondary signal for AT1 fate. In early postnatal and juvenile life it prevents AT2 reprogramming to AT1 fate, thereby consolidating the AT2 fate decision, whereas for the rest of life it serves as a survival signal, continuously suppressing AT2 apoptosis likely through Akt. AT2 cells also require Fgfr2 during alveolar repair ^32, 36^. Although there has long been evidence of Fgfr2 activities in the lung beyond its well-described role in airway branching^14, 18, 31–33, 36, 49, 50^, its multiple and changing roles, the difficulty in conditionally, rapidly and completely deleting pathway genes, and the rapid and robust adult regenerative response have obscured its critical, continuous, and ubiquitous functions in the selection and lifelong maintenance of AT2 cells.

An important future goal will be to molecularly elucidate how the FGF signaling pathway selects and maintains AT2 cells, and how it results in distinct cellular outcomes at different stages in lung development. The same ligand (*Fgf7, Fgf10*) and receptor (*Fgfr2*) genes are deployed throughout life, so presumably changes in the downstream signal transduction and effector pathway or in parallel pathways account for the stage-specific effects of Fgfr signaling.

Such a model is operative in Drosophila respiratory system development where the initial round of Fgfr signaling induces changes in expression of downstream effectors that alter the response in the next signaling round^51^. Appealing candidates for the differential apoptotic response of juvenile vs adult AT2 cells are anti-apoptotic BH3 proteins regulated by Akt activity^43, 52^, several of which (*Bag1, Mcl1, Bcl2l1*) are transiently upregulated in perinatal AT2 cells (D.G.B., unpublished observations) so could buffer them against apoptosis. Other possible stage-specific effectors include ETV4 and ETV5, related transcription factors induced by Fgf7 and Fgf10 at bud tips during airway branching^53^ that regulate bud size, growth, and number^54^ but whose expression restricts such that ETV5 becomes specifically expressed in the AT2 lineage^11^, where it is required to maintain expression of AT2-selective genes in cultured cells as well as the adult mouse lung^55^.

The Fgfr2 pathway remains active in AT2 cells and sustains them throughout life (Fig. 7J). Why would a developmental signaling pathway remain on for months or even years after its developmental role has been completed, simply to keep the selected cell alive? The considerable energy expenditure involved in lifelong signaling suggests some substantial benefit of active AT2 maintenance. Perhaps it allows rapid conversion to a progenitor state and other fates following alveolar injury to quickly restore gas exchange function, akin to the rapid transdifferentiation of club into ciliated airway cells when Notch signaling is blocked^47^. Or perhaps it prevents initiation of lung adenocarcinoma, the leading cancer killer, when an AT2 cell divides and a daughter loses contact with the underlying stromal cells that express the Fgf7 and Fgf10 survival signals (Fig. 7K). Whatever the reason, the pathway plays an important role in human lung health because genetic studies identify *Fgf7* as a susceptibility locus for the common and devastating disease COPD (chronic obstructive lung disease)^56^.

Our results show that loss or inhibition of Fgfr2 signaling has dire consequences for the alveolus and gas exchange, though they may not become apparent immediately because of the robust regenerative response activated by AT2 cell death. However, widespread targeting or loss of AT2 cells, as may occur in SARS, MERS, and COVID-19 coronavirus infections^6–8^, causes acute and severe alveolar injury and can be rapidly fatal. Pharmacologic modulation of Fgfr2 signaling could be used to support alveolar health during acute injuries like these, helping sustain AT2 cells as a virus or toxin destroys them^6–8^, and perhaps similarly in chronic diseases such as COPD/emphysema and pulmonary fibrosis^57^. Modulators could also be used to create alveolar cells for in vitro studies or cell therapies for these diseases.

## Acknowledgments

The authors thank members of the Krasnow laboratory for helpful discussions and critical reading of the manuscript. This work was supported by R00HL127267 (D.G.B), NHLBI U01 Progenitor Cell Biology Consortium grant and the Howard Hughes Medical Institute (M.A.K.), Ludwig Cancer Center (T.J.D. and M.A.K.), Chan Zuckerberg Institute (T.J.D. and M.A.K.), NIH T32HD007249 (D.G.B.), and NHLBI 5R01HL14254902 and NIH 5UG3HL14562302 (T.J.D.). D.G.B. is the Mark and Catherine Winkler Assistant Professor of Cell and Developmental Biology, T.J.D. is the Woods Family Endowed Faculty Scholar in Pediatric Translational Medicine of the Stanford Child Health Research Institute, and M. A. K. is the Paul and Mildred Berg Professor at Stanford University and an Investigator of the Howard Hughes Medical Institute.

## Author Contributions

DGB, TJD, and MAK conceived the study and designed the experiments. DGB and TJD performed the experiments. DGB, TJD, and MAK analyzed and interpreted the data. DGB, EG, and AG performed immunostaining in the embryonic mouse lung. AD and EG performed tracheal instillations and AD performed multiplex in situ hybridization. DGB, TJD, and MAK wrote the paper.

## Methods

### Mouse strains

Timed-pregnant C57BL/6J females (abbreviated B6; Jackson Laboratories) were used for all embryonic time points, with gestational age verified by crown–rump length. For studies of adult wild type lungs, B6 males and females were used. Mosaic labeling and deletion studies were conducted by Cre recombinase expression using gene targeted alleles *BAC-Nkx2.1-Cre* ^61^, *Lyz2* (also called *LysM*)*-Cre* ^62^, and *SftpC-Cre-ERT2-rtTA* ^63^. Cre-dependent target genes were the conditional "floxed" *Fgfr2^fl/fl^* for removal of *Fgfr2* ^64^, Cre reporter *Rosa26-mTmG,* which expresses membrane-targeted tdTomato (mT) in all tissues and mGFP following Cre-mediated recombination ^65^, and the Flp-dependent *Rosa26-MASTR* allele that constituitively expresses Cre recombinase following Flp recombination ^37^. For initiating recombination with *SftpC-Cre-ERT2*, intraperitoneal injections of 3 mg tamoxifen were administered twice a week for two weeks, except where noted. Genomic DNA was extracted from tails by Proteinase K (Sigma) digestion and genotyping performed by polymerase chain reaction (PCR) using published primer sets. Mice were housed in filtered cages and all experiments were performed in accordance with approved Institutional Animal Care and Use Committee protocols at Stanford University.

### Lung isolation and processing

For prenatal time points, individual embryos were staged by fetal crown-rump length before sacrifice and lungs removed en bloc. For postnatal time points, mice were euthanized by carbon dioxide inhalation and dissected to both expose the lungs and allow for exsanguination via the abdominal aorta. Phosphate buffered saline (PBS; Ca^2+^-and Mg^2+^-free, pH 7.4) was gently perfused into the right ventricle of the heart by syringe with 21 gauge needle until the lungs appeared white. For postnatal time points at or after 8 days, the trachea was then cannulated with a blunt needle and the lungs gently inflated to full capacity with molten low melting point agarose (Sigma, 2% in PBS). For all time points, lungs were then separated by lobe and fixed in either Zinc Formalin, 4% PFA, or Dent’s fixative at 4 °C. For immunostaining, a vibrating microtome (Leica) was used to generate embryonic (200 µm) or adult (450 µm) tissue sections of uniform thickness. For in situ hybridization, lobes were submerged in OCT (Tissue Tek) in an embedding mold and frozen on dry ice. Tissue sections (10 µm) were obtained using a Leica CM3050S cryostat, collected on glass slides, and subsequently processed.

### Immunostaining

Immunohistochemistry was performed as previously described^66^ using primary antibodies against the following epitopes (used at 1:500 dilution unless otherwise noted): pro-SftpC (rabbit, Chemicon AB3786), RAGE (rat, R&D MAB1179), E-cadherin (rat, Life Technologies ECCD-2), Podoplanin (hamster, DSHB 8.1.1), Mucin 1 (hamster, Thermo Scientific HM1630 and rabbit, Novus NB120-15481), Ki67 (rat, DAKO M7249), Fgfr2 (rabbit, SCBT SC-122), cleaved Caspase 3 (rabbit, Novus NB100-56708), GFP (chicken, Abcam ab13970), Fgfr2iiib-Fc (human, R&D Systems) and Phospho-p44/42 MAPK (rabbit, CST D13.14.4E). Primary antibodies were subsequently detected using Alexa Fluor-conjugated secondary antibodies (Life Technologies) unless noted otherwise and incubated in Vectashield with DAPI (5 µg/ml, Vector labs). EdU was detected using the standard Click-iT reaction (ThermoFisher). Images were acquired using a laser-scanning confocal microscope (Zeiss LSM 780) and subsequently processed using ImageJ.

### In situ hybridization

Two methods of multiplexed single molecule fluorescence in situ hybridization (smFISH) of mRNAs was performed. To simultaneously detect *Sftpc*, *Fgf7*, and *Fgf10* RNAs, RNAscope multiplex assay V1 kit (ACD) was used as described^11^. To visualize *Sftpc* and *Fgfr2* RNAs, proximity ligation in situ hybridization (PLISH) was used^67^. Briefly, 20µm thick sections were cut from OCT-embedded, cryopreserved tissue and hybridized with multiple anti-sense probe pairs that hybridize to the target transcript. Probe pairs that targeted each gene share a common barcode that is unique and complementary to circle and bridge constructs. The circle and bridge constructs undergo proximity ligation to form a closed circle that undergoes rolling circle amplification. Detection oligonucleotides conjugated to fluorophores anneal to the rolling circle amplification product, generating discrete puncta for each transcript. The following sets of primer pairs were used to detect transcripts of the indicated genes:

*Sftpc*

5’ TCGTACGTCTAACTTACGTCGTTATGTGCGGTTTCTACCGACC3’

5’GGTCTTTCCTGTCCCGCTTATACGTCGAGTTGAAGAACAACCTG3’

5’TCGTACGTCTAACTTACGTCGTTATGTTTATTCTTTTGTGATAGGATCCC3’

5’TTGTTTTCCAATCAGGCTGCTTATACGTCGAGTTGAAGAACAACCTG3’

*Fgfr2*

5’TTAGTAGGCGAACTTACGTCGTTATGTCACCAGCGGGGTGTTGGAG3’

5’TTAGTAGGCGAACTTACGTCGTTATGTCATGTTTTAACACTGCCGTTTATGTGTGG3’

5’TTAGTAGGCGAACTTACGTCGTTATGAGACATGCTCATCGGACAGCAGAGT3’

5’TGTTGAGGACAGACGCGTTGTTATCCTTATACGTCGAGTTGAACATAAGTGCG3’

5’TGTTTGGGGACAGGAAGACACATTCACTTATACGTCGAGTTGAACATAAGTGCG3’

5’GTTTCTTGAAACATGGGCATTAGGGTGTCTTTATACGTCGAGTTGAACATAAGTGCG3’

Expression was detected by confocal fluorescence microscopy as ∼0.5 µm fluorescent puncta.

### Cell isolation and culture

Adult AT2 cells and embryonic (e16.5) bipotent alveolar epithelial progenitors were purified as previously described^11^. Adult 2-month-old mice were euthanized by administration of CO_2_. For e16.5, embryos were removed from the mother and their lungs were isolated en bloc without perfusion and pooled by litter (five to seven embryos) for further processing. Lungs were microdissected to remove the proximal lung tissue, leaving only the distal (alveolar region) tissue. Distal e16.5 lung cells were dissociated in dispase (BD Biosciences) and triturated with glass Pasteur pipettes until a single-cell suspension was attained. For adult lungs, the vasculature was perfused through the right ventricle with 37°C media (DMEM/F12, Life Technologies). The trachea was punctured and lungs inflated with digestion buffer (DMEM/F12 containing elastase (1U/ml, Worthington) and dextran (10%, Sigma)) for 20 minutes at 37°C. Digested lungs were minced with a razor blade into 1mm^3^ fragments, suspended in 5ml of digestion buffer containing DNase I (0.33U/ml; Roche), incubated with frequent agitation at 37°C for 45 minutes, and triturated briefly with a 5-ml pipette.

To deplete red blood cells, an equal volume of DMEM/F12 supplemented with 10% FBS and penicillin–streptomycin (1U/ml; Thermo Scientific) was added to the lung single-cell suspensions before filtering through 100µm mesh (Fisher) and centrifuging at 400g for 10 min. Pelleted cells were resuspended in red blood cell lysis buffer (BD Biosciences), incubated for 2 minutes, passed through a 40µm mesh filter (Fisher), centrifuged at 400g for 10 minutes and then resuspended in MACS buffer (2mM EDTA, 0.5% BSA in PBS, filtered and degassed) for purification.

From the resultant single cell suspensions of distal e16.5 lung, bipotent progenitors were isolated by MACS using MS columns (Miltenyi Biotec) according to the vendor protocol. Before column loading, suspensions were passed through a 35µm cell strainer (BD Biosciences). Other cell types were first depleted with antibodies against CD31, CD45 and Pdgfrα (Miltenyi Biotec), then bipotent progenitors were positively selected using a biotinylated EpCAM antibody (clone G8.8, eBiosciences) and streptavidin-conjugated magnetic beads (Miltenyi Biotec). This generates a >95% pure preparation of e16.5 bipotent alveolar progenitors (Sftpc^+^ Rage^+^ cells). AT2 cells were isolated from the single cell suspensions of adult lungs in the same way, generating a >90% pure preparation of AT2 cells. To enrich for adult fibroblasts, MACS depletion was conducted as above using antibodies against CD31, CD45, and EpCAM.

For culturing bipotent progenitors, cell density was calculated using a hemocytometer. A density of 100,000 cells per well was used for culture in 8-well #1 coverglass chambers (Labtek) precoated with growth factor reduced Matrigel (80µl, BD Biosciences) for 30 minutes at 37°C. Cells were supplemented with Fgf7 (50ng/ml, R&D Systems), Fgf10 (100ng/ml, R&D Systems), FIIN-1 (20nM, Tocris), Fgfr2iiib-Fc (1µg/ml, R&D Systems), CK666 (40µM, R&D), and HSPG (100ng/ml, Sigma) to the indicated concentrations in DMEM/F12. Cells were maintained at 37°C in 400µl of DMEM/F12 with media changes every other day in a 5% CO_2_/air incubator typically for four days, except where indicated otherwise.

### Live imaging of alveolar progenitors in culture

Time lapse microscopy of cultured alveolospheres was conducted using an inverted LSM 880 equipped with environmental control to maintain 37°C and 5% CO_2_. Isolated bipotent progenitors were plated and cultured for ∼12 hours before confocal microscopy. Brightfield z-stacks (20x multi-immersion objective, NA 1.33) were acquired every 15 minutes for at least 3 days. Lumenal area, measured at the center z-position of each organoid, was used as an indicator of stretch, as commonly implemented in forskolin-induced swelling experiments^68^. Budding status of a cell was determined using the published criteria^18^ of basal extrusion with substantial reduction of lumenal cell surface.

### Single-cell RNA sequencing

Processing, scRNAseq, and analysis of e18.5 and adult mouse lung cells was conducted as previously described with minor alterations to enrich for the respective populations^11^. Briefly, single-cell suspensions sorted by MACS as CD45^-^ CD31^-^ EpCAM^-^ (for mesenchymal cells) or CD45^-^ CD31^+^ EpCAM^-^ (for endothelial cells) were subsequently incubated with a viability stain (Sytox Blue, Life Technologies) for 15 minutes and loaded on a medium-sized (10–17µm cell diameter) microfluidic RNA-seq chip (Fluidigm) using the Fluidigm C1 system. Cells were loaded at a concentration of 300–500 cells/ml and imaged by phase-contrast and fluorescence microscopy to assess the number and viability of cells per capture site. Only single, live cells were included in the analysis. For scRNAseq experiments, cDNAs were prepared on chip using the SMARTer Ultra Low RNA kit for Illumina (Clontech). ERCC (External RNA Controls Consortium) RNAspike-in Mix (Ambion, Life Technologies) was added to the lysis reaction and processed in parallel to cellular messenger RNA. Sequencing libraries were constructed (Illumina Nextera XT DNA Sample Preparation kit), and single-cell libraries pooled and sequenced 100 base pairs (bp) paired-end to a depth of 2 to 6 x10^6^ reads. Sequencing data were processed as described^11^, and transcript levels quantified as fragments per kilobase of transcript per million mapped reads (FPKM) generated by TopHat/Cufflinks. Cells not expressing (FPKM < 1) three of four housekeeping genes (*Actb*, *Gapdh*, *Ubc*, *Ppia*), or expressing them less than three standard deviations below the mean, were removed from analysis. Fibroblasts were identified as cells expressing canonical fibroblast markers *Mgp*, *Col1a1*, and *Col1a2* that also lacked expression of canonical markers of the major airway or alveolar epithelial cell types. Subsequent analysis including hierarchical clustering was performed as described^11^. To determine isoform expression of *Fgfr2*, STAR was used to align and count reads specific to either exon IIIb or IIIc To determine receptor genes restrictively expressed along either AT1 or AT2 lineage, receptor genes were first screened for ones detectably expressed in at least 6 cells, then Welch’s t-test was used to identify receptor genes with at least two-fold higher expression in each respective lineage (p ≤ 0.01).

For the analysis of cell type specific gene expression in published scRNAseq datasets, the processed mRNA counts for each cell were used from developing (SE119228)^10^ and adult mouse lung (GSE109774)^60^ datasets. Expression matrices were analyzed via R using the Seurat package. Clusters of cells with similar expression profiles were identified via shared nearest neighbor analysis using the Louvain algorithm, clusters were visualized in UMAP plots, and expression of specific genes was represented in heatmaps, violin plots, and dot plots generated in Seurat.

### Mosaic and inducible deletion of *Fgfr2*

To mosaically modulate Fgfr2 activity in culture, a lentiviral approach was developed using the Lenti-X Tet-On 3G Inducible Expression System (Clontech). The ORF encoding Fgfr2iiib (Origene; MC221076), the IRES sequence from pLVX-IRES-Puro (Clontech; 632186), and sfGFP (superfolder GFP) coding sequence from pBAD-sfGFP (Addgene; 54519) were PCR-amplified with adaptor sequences (IDT) and assembled (NEB; E5520) into the lentiviral backbone pTetOne (Takara; 631018) linearized by EcoRI/BamHI digestion (NEB) to generate pTetOne-Fgfr2-IRES-sfGFP (Fig. 3A). To generate a dominant negative form of Fgfr2iiib, a truncated version of Fgfr2iiib lacking the tyrosine kinase domain^69^ was also PCR-amplified with adapter sequences (as above) to assemble pTetOne-Fgfr2^DN^-IRES-sfGFP. Lentiviruses ("Lenti-Fgfr2", "Lenti-Fgfr2^DN^") were produced from these plasmids following the manufacturer’s instructions for transfection, lentiviral concentration, and titer estimation (Lenti-X Expression System, Clontech). An AAV virus that constitutively expresses eGFP from CMV promoter (Addgene, 105545-AAV9) was used as a cell labeling control (Supplementary Fig. 4B). Purified bipotent progenitors plated on Matrigel in standard medium as above were treated with cationic polymer polybrene (8ng/µl, EMD Millipore, to enhance viral infection) 10 minutes before viral addition. After 24 hours of infection at 37°C in media containing doxycycline (100ng/ml, Sigma) to induce expression of the Fgfr2iiib genes, media was replaced with media containing doxycycline and Fgf7 (50ng/ml). Four days later, cultured cells were fixed and analyzed by immunostaining.

For in vivo mosaic labeling and deletion of the floxed *Fgfr2* allele in bipotent progenitors, *Tg^Nkx^*^2^.^1^*^-Cre^*; *Rosa26*^mTmG/mTmG^ mice were used. For analogous experiments in AT2 cells, *Lyz2^Cre^*; *Rosa26*^mTmG/mTmG^ and *S*ftpcCre-ERT2-rtTA; *Rosa26*^mTmG/mTmG^ were bred and induced by intraperitoneal injection of tamoxifen. At time points and conditions indicated, lineage-labeled (mGFP+) and unlabeled (mGFP-) cell types (BP, AT2, or AT1, as indicated) identified by marker expression were counted from large alveolar fields (25x objective, ∼500µm x 500µm x 100µm).

To mosaically but efficiently delete *Fgfr2* in adult AT2 cells, we used a recombinant AAV described below to infect and express Flp recombinase mosaically in AT2 cells, and the Flp-dependent *Rosa26-MASTR* allele to constituitively express Cre recombinase^37^ to delete the conditional *Fgfr2^fl/fl^* allele. The pAAV-EF1α-Flpo plasmid^70^ was modified using the assembly scheme described above (NEB; E5520) to replace the EF1α promoter (removed by MluI/BamHI restriction digestion) with a previously defined^38^ 320 bp minimal promoter element of *Sftpc* that was chemically synthesized with added adapter sequences (IDT) (Fig. 6C). Recombinant AAV-Sftpc-Flpo virus was prepared by the HHMI Janelia Farm Viral Tools facility (2x10^14^ GC/ml) and endotracheally instilled (1µl of viral solution diluted in 50µl PBS) into the lungs of mice harboring *Rosa26^MASTR/mTmG^* and either heterozygous or homozygous for the floxed *Fgfr2* allele. Efficiency of *Fgfr2* deletion was assessed by immunostaining for Fgfr2. For adult time points, the density of AT2 cells was calculated and expressed as the number of AT2 cells per 100µm x 100µm x 100µm region of alveolar tissue. To determine AT2 proliferation, EdU (1mg/ml, Cedar Lane) was administered in the drinking water for the indicated period.

### Pharmacologic inhibition of Fgfr2 and activation of Akt

To rapidly and persistently inhibit Fgfr2 signaling, the irreversible Fgfr2 inhibitor FIIN-1 (14.75 mg/kg, Tocris)^71–73^ or recombinant Fgfr2iiib-Fc (100µg/30g mouse, R&D Systems) was used. Mice at least 2 months (and up to over 1 year) of age carrying the SftpcCre-ERT2-rtTA; Rosa26^mTmG/mTmG^ alleles were administered tamoxifen (3mg/30g mouse, Sigma) via intraperitoneal injection to label >95% of AT2 cells. After 1 week, the mice were anesthetized with isoflurane and administered by endotracheal instillation either vehicle alone (PBS with biotinylated WGA), or vehicle with the inhibitor. To test the effect of Akt signaling, the small molecule Akt activator SC79 (Tocris) was administered (0.04 mg drug per g mouse) via intraperitoneal injection 30 minutes prior to Fgfr2 inhibition. Lungs were isolated at the indicated time points and areas of high exposure to inhibitor were identified by WGA immunostaining.

### Statistical analysis

Data analysis and statistical tests were performed with R or GraphPad Prism software. Replicate experiments were all biological replicates with different animals, and quantitative values are presented as mean ±S.D. unless indicated otherwise. Student’s t-tests were two-sided. No statistical method was used to predetermine sample size, and data distribution was tested for normality prior to statistical analysis and plotting. Both male and female animals were used in experiments, and subjects were age- and gender-matched in biological replicates and in comparisons of different groups. Post-hoc power analysis was done (Supplemental Table 1).

## List of Supplementary Materials

### Legends of Supplementary Figures

**Supplementary Figure 1.**
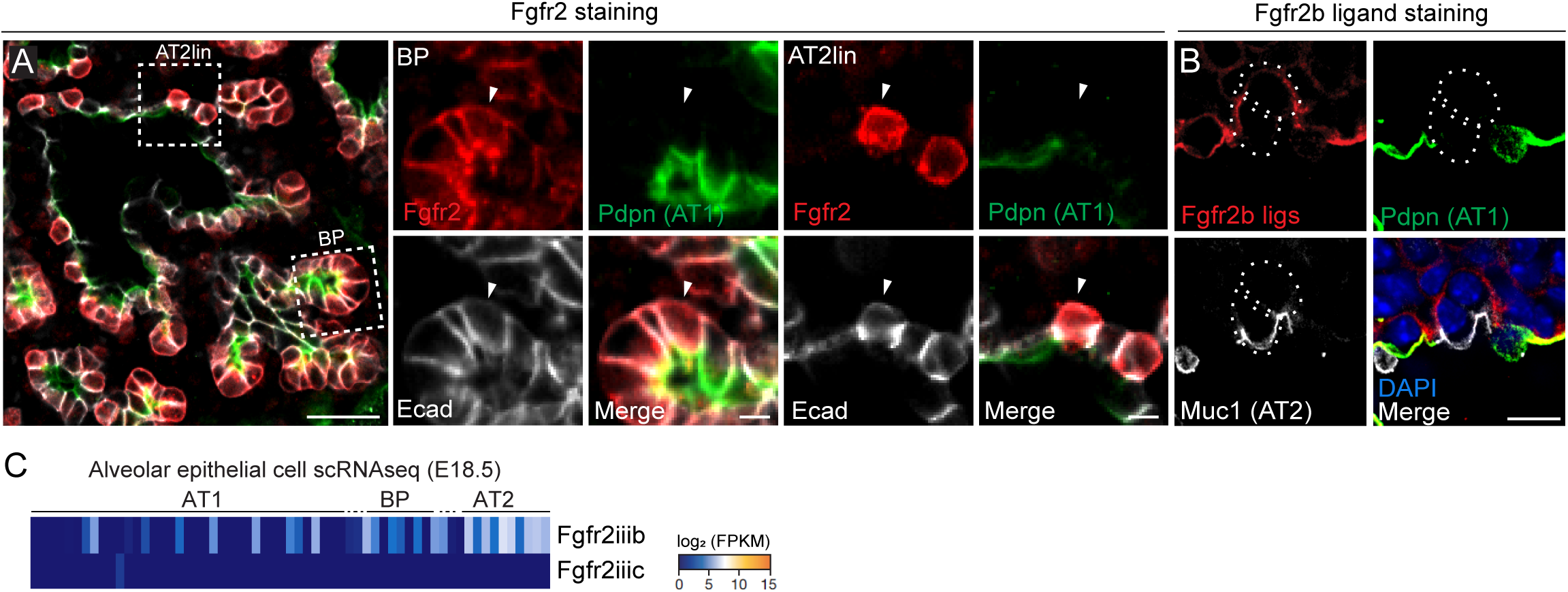
Additional examples of expression of Fgfr2 and its ligands during alveolar differentiation. (A) e17.5 lung immunostained for Fgfr2, AT1 marker podoplanin (Pdpn), and epithelial marker E-cadherin (E-cad) as in Fig. 1C. Boxed regions, close-ups and split channels at right of bipotent progenitor (BP) and developing AT2 lineage cell (AT2 lin). Note developing AT2 lineage cell (arrowhead) has lost expression of Pdpn so is more advanced in development than one shown in Fig. 1C that still expresses it. Scale bars, 50µm (left panel) and 10µm (right panels). (B) e17.5 lung stained with the Fgfr2 (isoform iiib) ligand-binding domain fused to human IgG1 domain as in Fig. 1D to show Fgfr2b ligands, and co-stained for Pdpn, and AT2 marker mucin1 (Muc1). Note two adjacent developing AT2 cells (dotted circles) and diffuse distribution of Fgfr2b ligands surrounding their basal surface and that of neighboring AT1 cells. Scale bar, 10µm. (C) scRNAseq heatmap of E18.5 alveolar epithelium as in Fig. 1A except Fgfr2 transcripts specific to Fgfr2iiib and Fgfr2iiic isoforms were mapped and calculated separately. Note Fgfr2iiib is almost exclusively expressed.

**Supplementary Figure 2.**
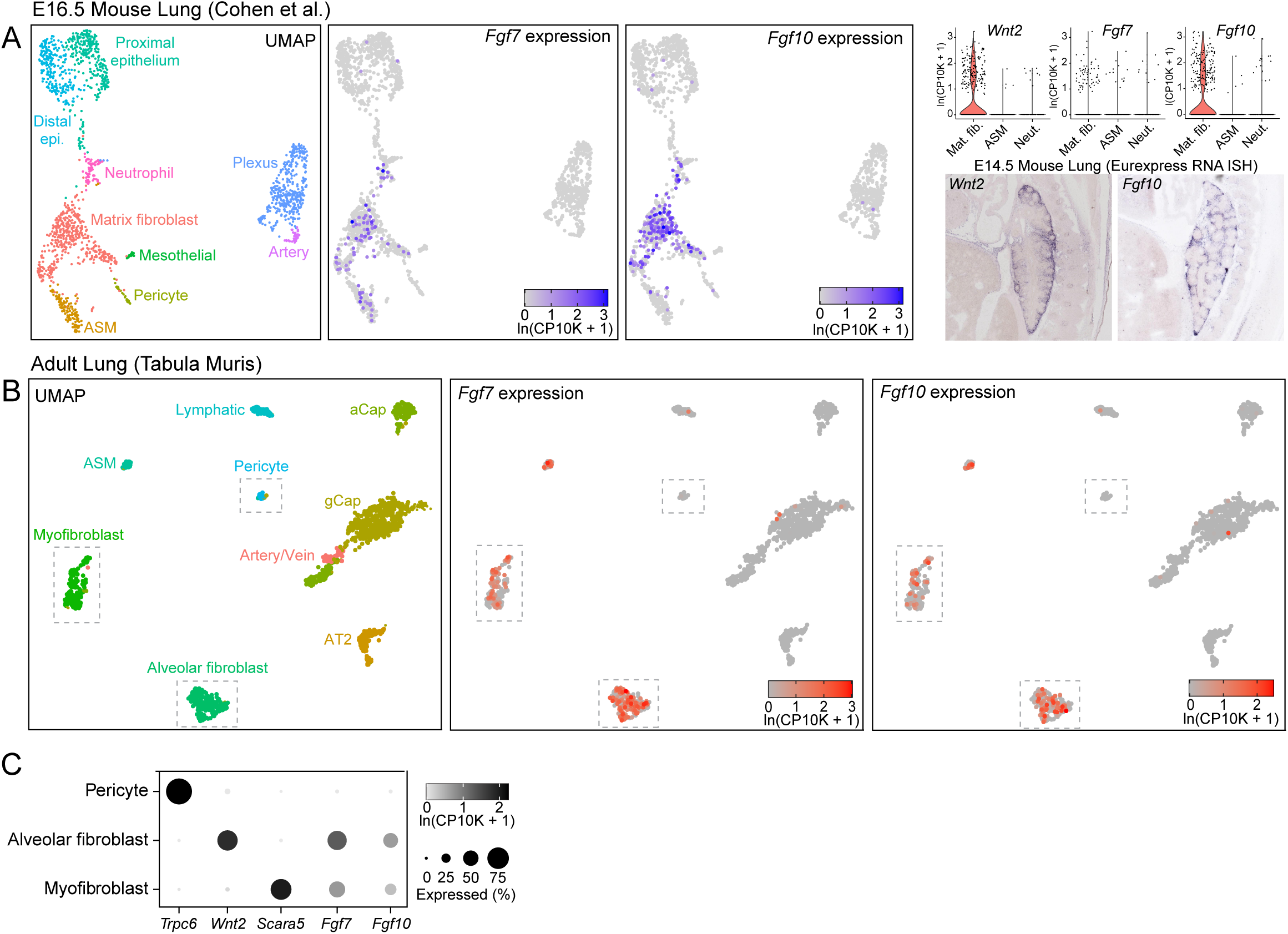
Identification by scRNAseq of the mesenchymal Fgf ligand source in the developing and adult mouse lung. (A) UMAP of transcriptomic profiles of e16.5 mouse lung cells^10^ clustered via Louvain algorithm and cell clusters identified as matrix fibroblast (*Mfap4, Wnt2*), airway smooth muscle (ASM, *Enpp2*), pericyte (*Gucy1a3*), mesothelial (*Wt1*), neutrophil (*Retnlg*), distal epithelial (*Sox9*), proximal epithelial (*Sox2*), capillary plexus (*Aplnr*), and arterial (*Gja5*) cell types by canonical marker gene expression^10, 11, 58, 59^ (Fib., fibroblast; Epi., epithelial). Gene expression values in heatmaps and violin plots were normalized, scaled and and log-transformed using Seurat (https://satijalab.org/seurat/) and given as ln(CP10K + 1)). Heatmaps (left) and violin plots (upper right) for *Fgf10* and *Fgf7* show both are expressed predominantly in matrix fibroblasts (Mat. fib., *Wnt2*-expressing) with sparse expression in airway smooth muscle (ASM) and neutrophils (Neut). (Lower right) RNA in situ hybridization of lung sections from e14.5 embryos (Eurexpress) show similar spatial expression patterns of *Wnt2* and *Fgf10*. (B) UMAP of transcriptomic profiles of adult mouse lung cells^60^ clustered via Louvain algorithm and cell clusters identified by canonical marker expression (ASM, *Acta2*; myofibroblast (Myofib), *Scara5*; alveolar fibroblast (Alv. Fib) , *Wnt2*; pericyte, *Trpc6*; lymphatic endothelium (Lymph), *Gja1*; aerocyte capillary (aCap), *Apln*; general capillary (gCap), *Aplnr*; artery/venous endothelium (Art/Vein), *Vwf*; AT2, *Sftpc*)^11, 58, 59^. Among the alveolar mesenchymal cell types (dashed boxes), note expression of both *Fgf7* and *Fgf10* in alveolar fibroblasts and, to a lesser extent, in myofibroblasts but not in pericytes. (C) Dotplot showing quantification (dot intensity, expression level; dot size, percent of cells with expression > 0) of *Fgf7* and *Fgf10* expression values from panel B along with selected cell type marker genes.

**Supplementary Figure 3.**
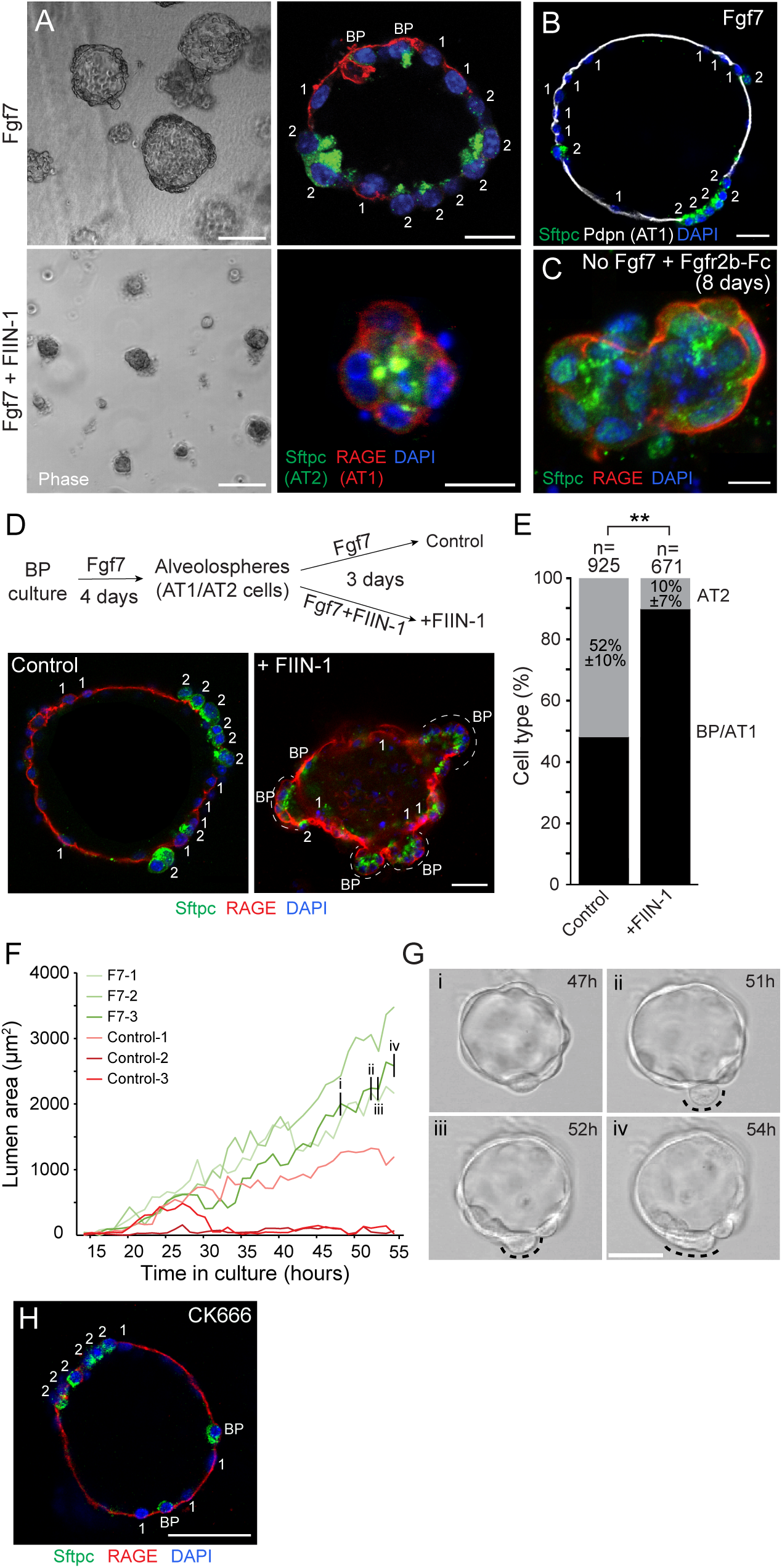
Live imaging of alveolosphere development in culture and effects of Fgfr and Arp2/3 inhibitors. (A) Phase images (left) and immunostains (right) of epithelial progenitors purified from tips of e16.5 lungs and cultured in Matrigel as in Fig. 2 for four days with Fgf7 alone (50 ng/ml added every two days; upper panels) or with Fgfr inhibitor FIIN-1 (10 nM; bottom panels). Note that Fgfr inhibition prevents bipotent progenitors (co-express Sftpc and RAGE) from forming alveolospheres (left) with differentiated AT1 and AT2 cells (right, same image as Fig. 2). BP, bipotent progenitor; 1, AT1 cell; 2, AT2 cell. Scale bars, 20µm. (B) Immunostaining of alveolospheres cultured with Fgf7 for 4 days and stained for the AT1-specific marker Podoplanin (white). Note expression is restricted to flat, Sftpcneg AT1 cells. Scale bar, 20µm. (C) Culture of e16.5 progenitors for 8 days in culture without FGF addition and with Fgfriiib-Fc (1µg/ml) added to block any FGF ligands provided by growth-factor-reduced Matrigel. Note progenitors remain undifferentiated bipotent progenitors (BP; Sftpc^+^ RAGE^+^) even after this extended culture. Scale bar, 10µm. (D) Alveolospheres generated by culturing progenitors for 4 days as in A and then cultured an additional three days in the continued presence of Fgf7 (50ng/ml) alone as control (left) or with added FIIN-1 (right, 10nM). Note loss of AT2 cells (Sftpc^+^ Rage^-^, labeled "2") and concomitant increase in AT1 cells (Sftpc^-^ RAGE^+^, labeled "1") and bipotent progenitors (Sftpc^+^ RAGE^+^, labeled "BP"). Scale bar, 20µm. (E) Quantification of panel D for AT2 cells (Sftpcpos Rage^neg^) and BP/AT1 cells (Sftpcpos RAGE^pos^ and Sftpcneg RAGE^pos^ cells). n, number of cells scored for each condition in experimental triplicate; **, p<0.001 (Student’s t-test). (F) Quantification of maximum luminal area in timelapse videomicroscopy recording of individual alveolospheres in control cultures without Fgf7 (red, Control-1, -2, -3) or with added Fgf7 (green, F7-1, F7-2, F7-3) as in A. Note continuous growth of all three alveolospheres in the cultures with Fgf7. (G) Frames at the indicated times from the alveolosphere F7-3 culture in panel F showing a transient cell budding (dashed arc) at hour 51, which soon regresses (hours 52, 54). Note luminal expansion has occurred (panel F) and AT1 flattening is already apparent at hour 47. Scale bar, 20µm. (H) Alveolosphere cultured in the presence of Fgf7 (50ng/ml) and the Arp2/3 inhibitor CK666 (40µM). AT2 and AT1 cells appear to develop normally, despite a reduction in transient budding events noted in live imaging (not shown). Scale bar, 20µm.

**Supplementary Figure 4.**
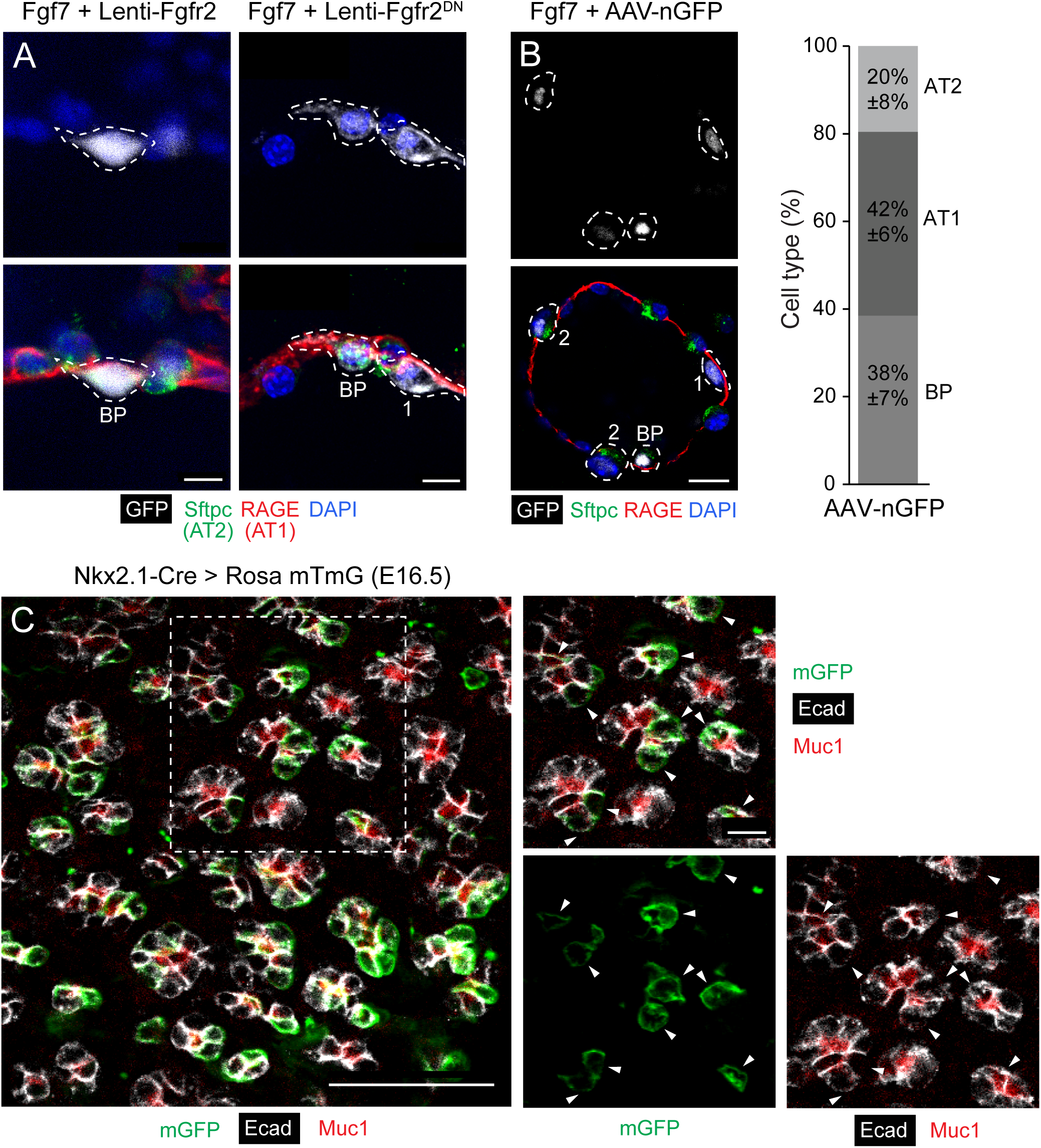
Mosaic cell labeling of alveolar cells in culture and in vivo. (A) Examples of the rare lentivirus-infected (GFP^pos^) cells (dashed outlines) from Fgf7-induced alveolospheres in Figure 3 that remained as a bipotent progenitor (BP, Sftpc+ RAGE+) despite infection with the indicated Fgfr2 lentivirus that caused nearly all infected cells to develop as AT2 cells (Lenti-Fgfr2, left panel) or AT1 cells (Lenti-Fgfr2^DN^, right panel; "1", Sftpc-Rage+) (see Figure 3). Scale bars, 10µm. (B) Control alveolosphere as in Figure 3 except treated instead with AAV-nGFP prior to Fgf7 induction, then immunostained for indicated markers to determine cell identity (BP, AT1, AT2, left panel) acquired during culturing. Scale bar, 20µm. Quantification of acquired identities of infected (GFP^pos^) cells (right panel, n > 50 GFP^pos^ cells scored in experimental triplicate) shows that progenitors infected with this control virus (lacking *Fgfr2*) have a similar chance of acquiring either AT1 (42 +/-6%) or AT2 (38 +/-7%) fate. (C) Distal epithelium of e16.5 lung from control *Nkx2.1-Cre; Rosa26-mTmG* mouse (similar to Fig. 4) immunostained for mGFP, E-cadherin and Muc1, which labels bipotent progenitors at this stage. Right panels show close up (with split channels as indicated) of boxed region in left panel. Note sporadic mGFP^pos^ cells (arrowheads) intermixed with mGFP^neg^ cells in epithelium, showing Nkx2.1-Cre activity in distal epithelium is highly mosaic rather than uniform. Scale bars 50µm, 10µm (insets).

**Supplementary Figure 5.**
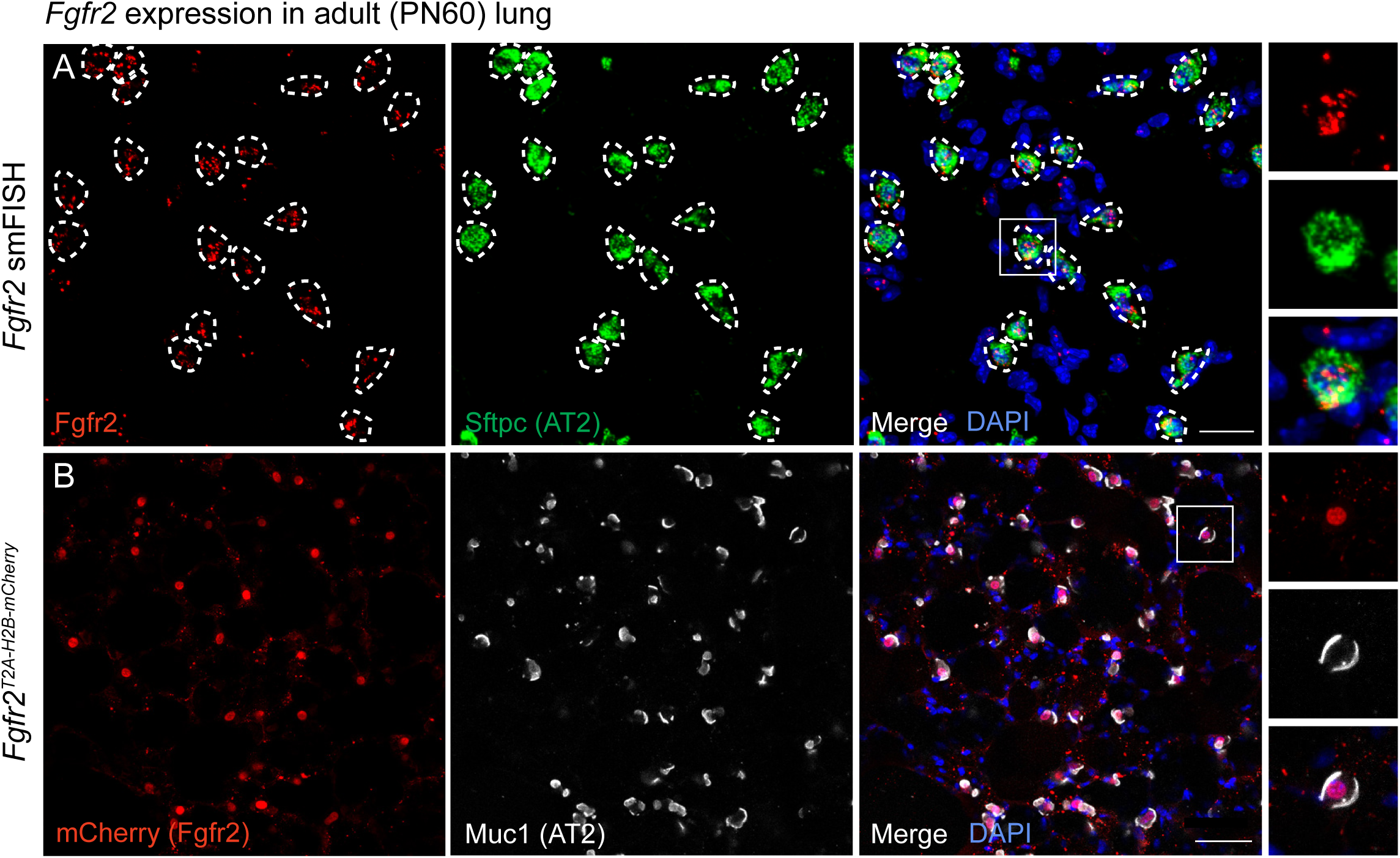
*Fgfr2* is selectively expressed in adult AT2 cells. (A) Single molecule fluorescence in situ hybridization (smFISH; PLISH) of alveolar region of adult (PN60) mouse lung for *Fgfr2* (red) and AT2 cell marker *Sftpc* (green). Right panels, close-up of boxed region. Note co-expression of *Fgfr2* and *Sftpc* mRNA indicating only AT2 cells express *Fgfr2*. Scale bar, 20µm. (B) Immunostain of alveolar region of adult (PN60) *Fgfr2^T2A-H2B-mCherry^* lung, carrying an *Fgfr2* knock-in reporter allele, immunostained for mCherry reporter and AT2 marker Muc1. Note selective expression of *Fgfr2* reporter in AT2 cells. Scale bar, 50µm.

**Supplementary Figure 6.**
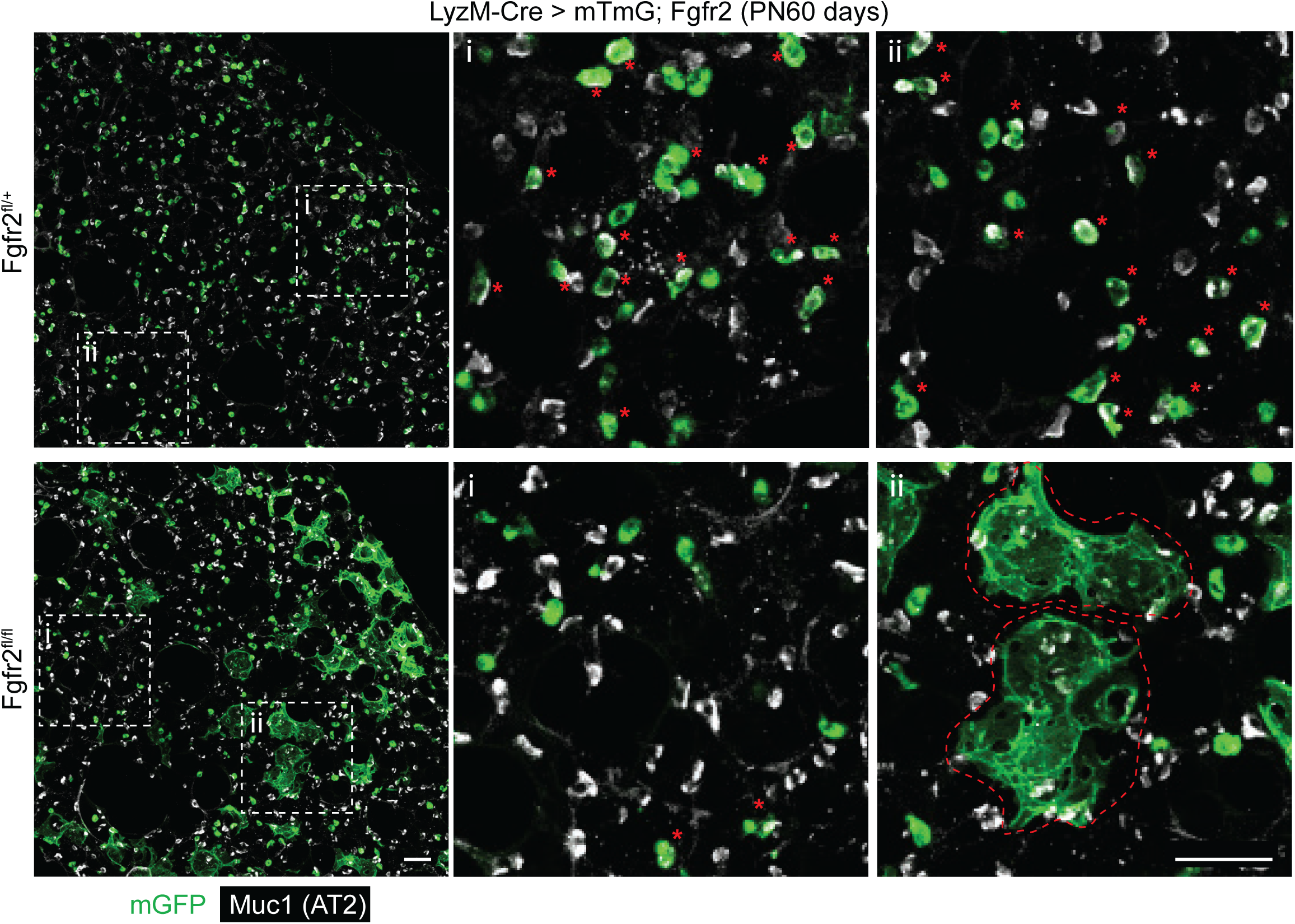
Large field of view of adult lungs following mosaic *Fgfr2* deletion in AT2 cells throughout life. Tiled maximal intensity projections showing large field of view of alveolar regions of adult (PN60) control Fgfr2^+^ *LyzM-Cre; Rosa26-mTmG; Fgfr2^fl/+^* (upper panels) and AT2 cell conditional deletion *LyzM-Cre; Rosa26-mTmG; Fgfr2^fl/fl^* (lower panels) mice (see Fig. 7A,B) immunostained for markers indicated. Although there are many regions in lung in lower panel (e.g. box i) where GFP lineage-labeled Muc1^pos^ AT2 cells (red asterisks) are depleted due to Fgfr2 deletion as in Fig. 7B (the other GFP lineage-labeled cells are alveolar macrophages, which lack Muc1), note there are also scattered mGFP lineage-labeled AT1 cells in this lung (box ii, lower panel; large cup-shaped cells, red dashed outlines), which presumably persist after their generation by *Fgfr2* loss from AT2 cells during the perinatal period. Scale bars, 50µm.

**Supplementary Figure 7.**
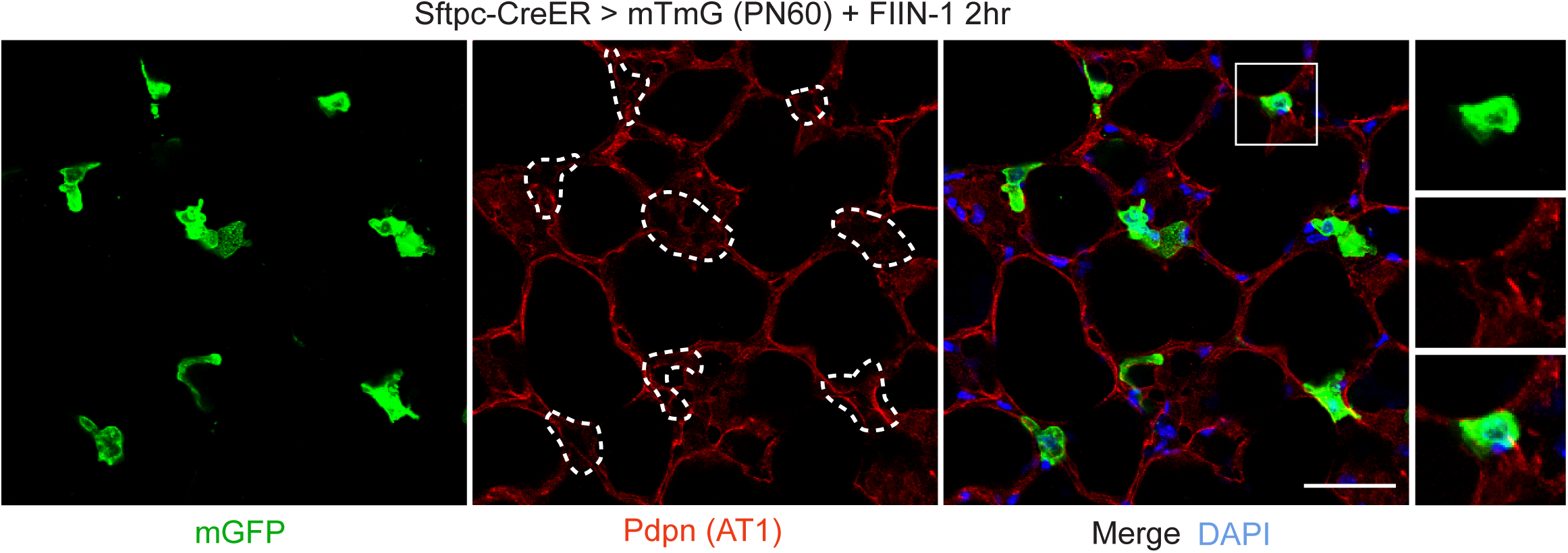
Fgfr inhibition in adult lung does not cause early reprogramming to AT1 fate. Alveolar region of adult (PN60) *Sftpc-CreER; Rosa26-mTmG* lung induced with tamoxifen to label AT2 cells with GFP, then one week later instilled with Fgfr inhibitor FIIN-1 as in Fig. 6C-F and stained 2 hours later for GFP and AT1 marker Podoplanin (Pdpn) with DAPI counterstain. Right panels, close-up of boxed region. Although many lineage-labeled (GFP^pos^) AT2 cells in exposed region are undergoing apoptosis at this time (see Fig. 6E,F), note that none of the lineage-labeled cells (dashed outlines; 0 of 398 GFP^pos^ cells scored in 3 experiments) showed evidence of reprogramming to AT1 fate (squamous morphology or expression of AT1 marker Pdpn). Scale bar, 50µm.

**Supplementary Figure 8.**
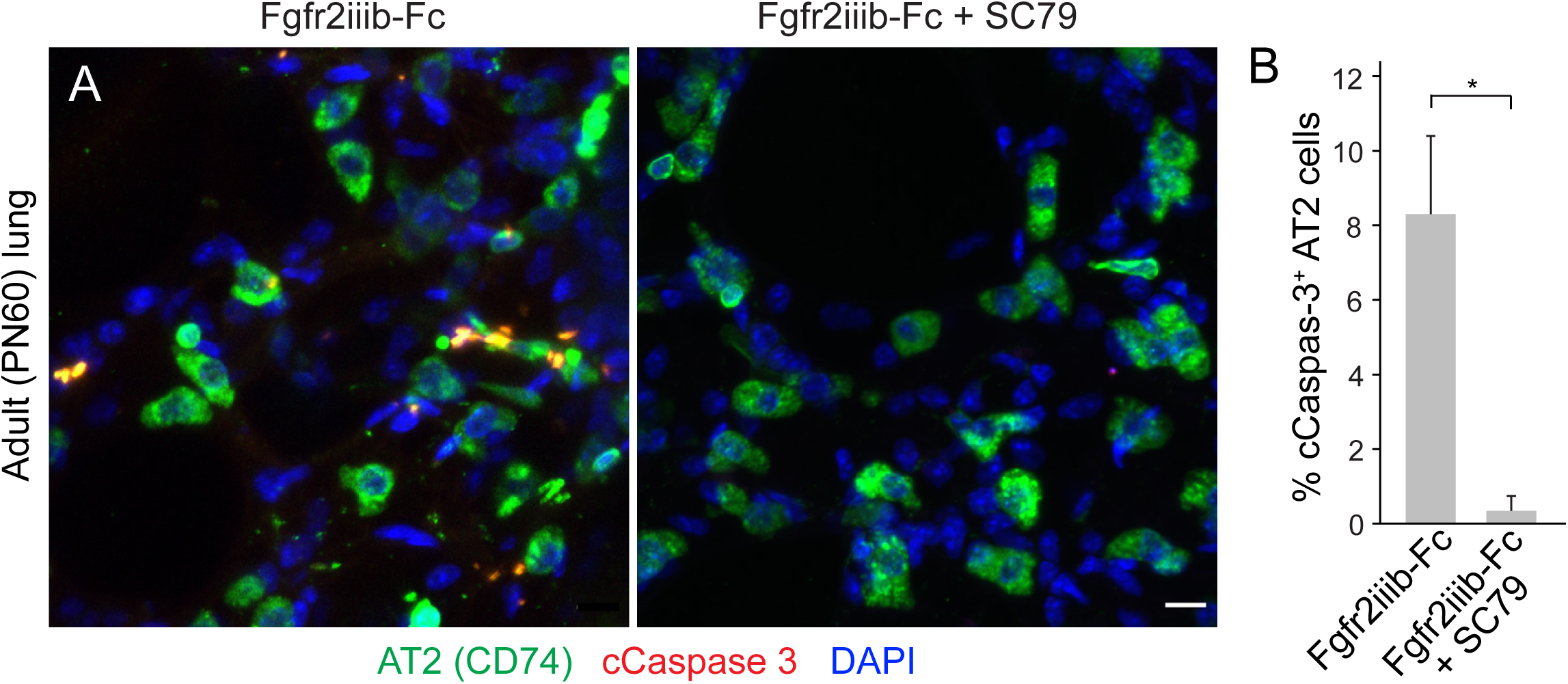
AT2 cell death following Fgfr2 inhibition is abrogated by Akt activation. (A) Close-ups of WGA-labeled alveolar regions from adult (PN60) lungs co-stained for AT2 marker CD74 and apoptotic marker cleaved Caspase 3 (cCaspase 3) and counterstained with DAPI 6 hours after the lungs were co-instilled with WGA-405 and Fgfr2iiib-Fc (as in Fig. 6O,P) without a pre-treatment as control (left) or with pre-treatment (intraperitoneal injection 30 min before WGA-405/Fgfr2iiib-Fc instillation) with Akt activator SC79 (0.04 mg/g mouse) (right). Note AT2 cell apoptosis (cCaspase 3 expression) following inhibition of Fgfr2 by Fgfr2iiib-FcAT2 (left) is prevented by Akt activation with SC79 (right). Scale bars, 10µm. (B) Quantification of A showing percent of AT2 cells that are apoptotic (cCaspase 3^+^) in control and SC79-pre-treated lungs. n>120 AT2 cells scored per condition in experimental triplicate; *, p-value <0.001 (Student’s t-test).

**Supplementary Figure 9.**
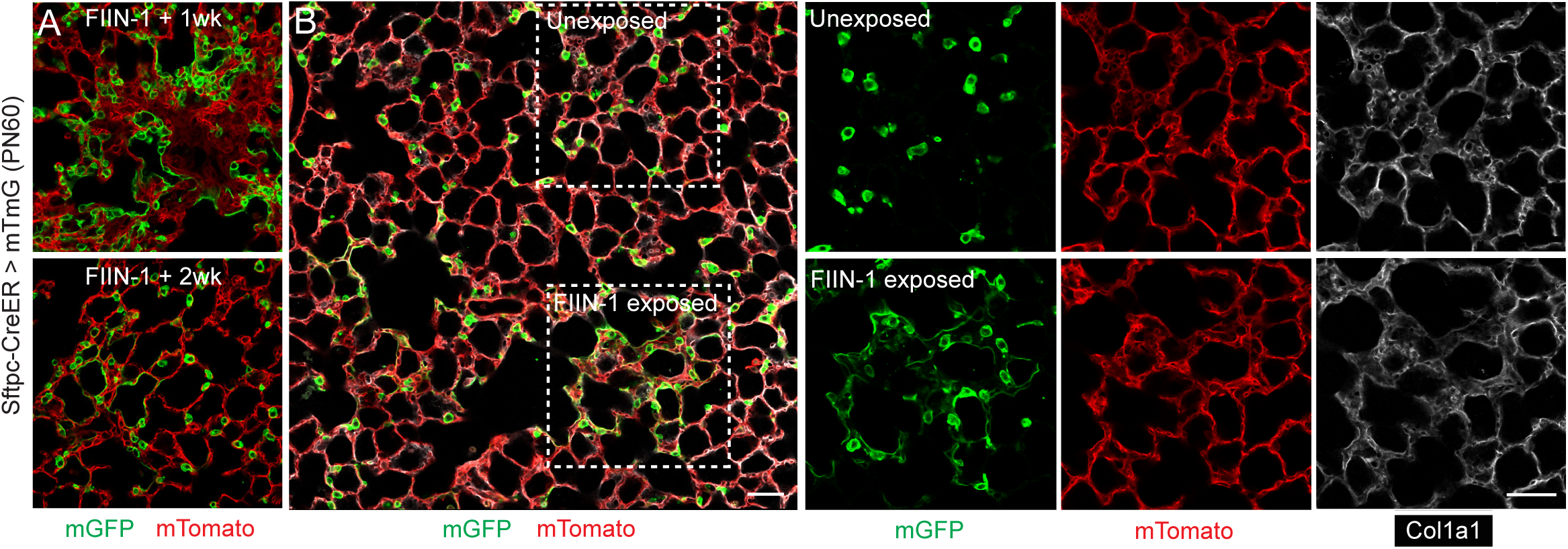
Alveolar restoration following acute Fgfr inhibition. (A) Alveolar regions of adult (PN60) *Sftpc-CreER; Rosa26-mTmG* lung induced with tamoxifen to GFP label AT2 cells, then 1 week later instilled with Fgfr inhibitor FIIN-1 as in Figure 6C-F and allowed to recover for one week (upper panel) or two weeks (lower panel). Note disorganization of alveolar structure at one week that resolves by two weeks. Scale bar, 50 µm. (B) FIIN-1-exposed alveolar region (upper box) and control unexposed region (lower box) following FIIN-1 instillation as above and stained two weeks later for AT2 lineage marker mGFP, mTomato (marking other cell types), and collagen a1 (Cola1). Note that the FIIN-1 exposed region has recovered and appears normal histologically with normal Col1a1 staining. Scale bars, 50µm.

### Supplementary Table

**Supplementary Table 1.** **Post-hoc power analysis for statistical tests.** Post-hoc power analysis of t-tests was done with standard values (alpha = 0.5) for a continuous endpoint, two independent sample study. Note power for each comparison was >0.99, indicating adequate sample size to determine a difference of this magnitude between the comparison groups. Power of goodness of fit test was done for chi-squared analysis (Fig. 3C). Power analysis was not done for Fig. 5I, which showed a non-normal data distribution; p-value shown was for Mann Whitney U test.

| Figure | Comparison | Power |
| --- | --- | --- |
| Figure 2 | Control versus Fgf7, Fgf10 | >0.99 |
| Figure 4B | Fgfr2 flox versus control | >0.99 |
| Figure 5H | Density AT2 versus Fgf+ | >0.99 |
| Figure 5P | % GFP+ AT2 cells, PN5 versus PN20 | >0.99 |
| Figure 5R | % GFP+ cells, PN5 versus PN20 | >0.99 |
| Figure 6F | fl/fl versus +/fl GFP+ AT2 | >0.99 |
| Figure 6J | fl/fl versus +/fl cCas3+ | >0.99 |
| Figure 6Q | Fgfr2iib-Fc versus veh. | >0.99 |
| Figure 7D | EdU+ AT2 cells in SftpC-CreER | >0.99 |
| Figure 7F | Ki67+ AT2 cells Fiin1 versus veh | >0.99 |
| Figure 3C | Control versus Fgfr2-WT, Fgfr2-DN<br>(Chi-Squared) | >0.99 |

| Figure | Comparison | p-value |
| --- | --- | --- |
| Figure 5I | Fgf+ vs AT2 (non-normal dist) | <0.001 |

